# The Effect of Ligands and Transducers on the Neurotensin Receptor 1 (NTS1) Conformational Ensemble

**DOI:** 10.1101/2021.12.08.471782

**Authors:** Austin D. Dixon, Asuka Inoue, Scott A. Robson, Kelly J. Culhane, Jonathan C. Trinidad, Sivaraj Sivaramakrishnan, Fabian Bumbak, Joshua J. Ziarek

## Abstract

Using a discrete, intracellular ^19^F-NMR probe on transmembrane helix 6 (TM6) of the Neurotensin receptor 1 (NTS1), we aim to understand how ligands and transducers modulate the receptor’s structural ensemble in solution. For apo NTS1, ^19^F-NMR spectra reveal an ensemble of at least three conformational substates (one inactive and two active-like) in equilibrium that exchange on the ms-s timescale. Dynamic NMR experiments reveal that these substates follow a linear three-site exchange process that is both thermodynamically and kinetically remodeled by orthosteric ligands. As previously observed in other GPCRs, the full agonist is insufficient to completely stabilize the active-like state. The inactive substate is abolished upon coupling to β-arrestin-1 or the C-terminal helix of Gα_q_, which comprises ⍰60% of the GPCR/G protein interface surface area. Whereas β-arrestin-1 exclusively selects for pre-existing active-like substates, the Gα_q_ peptide induces a new substate. Both transducer molecules promote substantial line-broadening of active-like states suggesting contributions from additional μs-ms exchange processes. Together, our study suggests i) the NTS1 allosteric activation mechanism may be alternatively dominated by induced fit or conformational selection depending on the coupled transducer, and ii) the available static structures do not represent the entire conformational ensemble observed in solution.

## INTRODUCTION

G protein-coupled receptors (GPCRs) serve as the primary hubs to relay changes in extracellular environments across the eukaryotic cell membrane.^1^ The more than 800 members of this protein superfamily share a conserved seven transmembrane helix (TM) bundle architecture that recognizes a large variety of ligands comprising small molecules, hormones, peptides, and photons.^2^ As such, it is no surprise they encompass over 30% of the drug market.^3^ Although atomic models are still relatively scarce compared to other protein classes, there are currently 121 unique receptor structures, or ~14% of the total GPCR superfamily.^4^ The difficulty of GPCR structural studies primarily reflects inherent protein instability and low recombinant expression. Through the use of detergent membrane mimetics and creative receptor engineering, the rate at which new receptor structures are determined has increased in recent years.^5^ These atomic models have revealed conserved, long-range allosteric activation networks that link the receptor orthosteric pocket to the intracellular bundle across the cell membrane. Most notably the DRY, PIF, CWxP, and NPxxY motifs serve as internal molecular “switches” of Class A GPCRs, connecting ligand-binding to downstream effector molecule complexation and activation events, spanning a distance of nearly 50 Å.^6^ Modeling of allosteric switches across numerous receptors has led to a putatively conserved structural activation profile.^7,8^

Neurotensin receptor 1 (NTS1) has quickly become one of the most well-characterized GPCRs with structures of the apo state, complexes with various pharmacological ligands, and ternary complexes with both the heterotrimeric G_i_ protein and β-arrestin-1 (βArr1) transducers.^9–17^ NTS1 is a Class A, β group receptor that is expressed throughout the central nervous system and gastrointestinal tract.^18^ Activation by its endogenous tridecapeptide ligand neurotensin (NT) mediates a variety of physiological processes including low blood pressure, high blood sugar, low body temperature, mood, and GI motility.^19^ It is also a long-standing therapeutic target for Parkinson’s disease, Schizophrenia, obesity, hypotension, psychostimulant substance use disorders, and cancer.^20^

Current atomic models derived from either X-ray crystallography or cryo-EM capture NTS1 in different stages of activation, mediated by bound ligands and transducer proteins. A hallmark of GPCR activation is the outward movement of transmembrane helix 6 (TM6) to accommodate G protein and arrestin complexation.^21^ In NTS1, ligand binding at the extracellular orthosteric pocket allosterically induces a ~13 Å lateral displacement at the intracellular tip of TM6.^10^ Ultimately, these models remain static. This has left a void in the literature detailing the NTS1 conformational ensemble and the pleiotropic effects ligands and transducers have on individual substates. This inspired us to pursue solution nuclear magnetic resonance (NMR) spectroscopy as a complementary approach to better characterize the allosteric activation mechanism in NTS1.

In this study, we ^19^F-label TM6 of a thermostabilized NTS1 construct solubilized in 2,2-didecylpropane-1,3-bis-β-D-maltopyranoside (LMNG) detergent micelles. Trifluoromethyl NMR probes are an optimal choice for site-selective isotopic labeling due to their low background signal and high spin-1/2 natural abundance.^25^ Their observed chemical shift value is dominated by solvent polarity and the local electronic environment, which makes them very sensitive to large conformational rearrangements observed in GPCRs. For example, as the intracellular tip of TM6 moves outward to accommodate transducer proteins, we anticipate an upfield chemical shift perturbation reflecting increased solvent exposure.^26^ NMR can provide both qualitative and quantitative information regarding the timescale of structural motions.^27^ While very fast rotation about the methyl axis averages any local fluctuations into a single peak, slower “biologically-relevant” motions on approximately the microsecond to milliseconds timescale affect both the resonance linewidth and chemical shift.^28^ As conformational exchange slows further into the millisecond to second regime, the averaged resonance will split into distinct peaks with characteristic linewidths and chemical shifts.^29^ Herein, we employ the G_q_ C-terminal α5-helix peptide and a pre-activated βArr1 to recapitulate responses to the heterotrimeric G_q_ protein and βArr1.^30,31^ Together, this enables us to develop a dynamic model of NTS1 activation in which ligands and transducers are allosterically coupled.^32^

## RESULTS

### Thermostabilized Neurotensin receptor 1 retains signaling activity

The well-characterized structure of NTS1 in a variety of pharmacologically-relevant states creates an ideal system for exploring the allosteric mechanisms of GPCR activation. Yet, wildtype NTS1 structural characterization remains challenging due to poor receptor stability following isolation from native membranes.^33^ All published NTS1 structures to date incorporate some combination of thermostabilizing mutations, lysozyme fusions, DARPin fusions, or conformationally-selective antibodies.^9,12,34,35^ Here, we employed a functional, thermostabilized rat (r)NTS1 variant (termed enNTS1) for solution NMR spectroscopy.^32^

To further validate enNTS1’s functional integrity, we performed a cell-based alkaline phosphatase (AP) reporter assay for G protein activation. Stimulation of Gα_q_ and Gα_12/13_ leads to ectodomain shedding of an AP-fused transforming growth factor-α (TGFα), which is then quantified using a colorimetric reporter.^22^ HEK293A cells were transfected with AP-TGFα and a NTS1 plasmid construct. A hexapeptide corresponding to NT residues 8-13 (NT8-13) is sufficient to generate a full agonist response in wildtype rNTS1;^36^ NT8-13 stimulates robust, concentration-dependent G protein-coupling to enNTS1 in the TGFα shedding assay, though with reduced efficacy compared to human (h)NTS1 (Figure 1A and Figure S1). Both enNTS1 and hNTS1 were equally expressed on the cell surface (Figure S1C). βArr1 recruitment was also measured using a NanoBiT enzyme complementation system.^23^ The large and small fragments of the split luciferase were fused to the N-terminus of βArr1 and the C-terminus of NTS1, respectively, and these constructs were expressed in HEK293A cells. As a negative control, we used the vasopressin V2 receptor (V2R) C-terminally fused with the small luciferase fragment. enNTS1 exhibited weak basal βArr1 recruitment that did not increase upon agonist addition (Figure 1B and Figure S1B). Nonetheless, addition of the βArr1-biased allosteric modulator (SBI-553) dose-dependently potentiates NT8-13-mediated βArr1 recruitment (Figure S1D).^37,38^ As SBI-553 alone is unable to substantially stimulate βArr1 recruitment to enNTS1 at the same concentration, we conclude that enNTS1 recruits using the same molecular mechanism as wildtype NTS1, although with reduced potency (Figure S1E).

**Figure 1.**
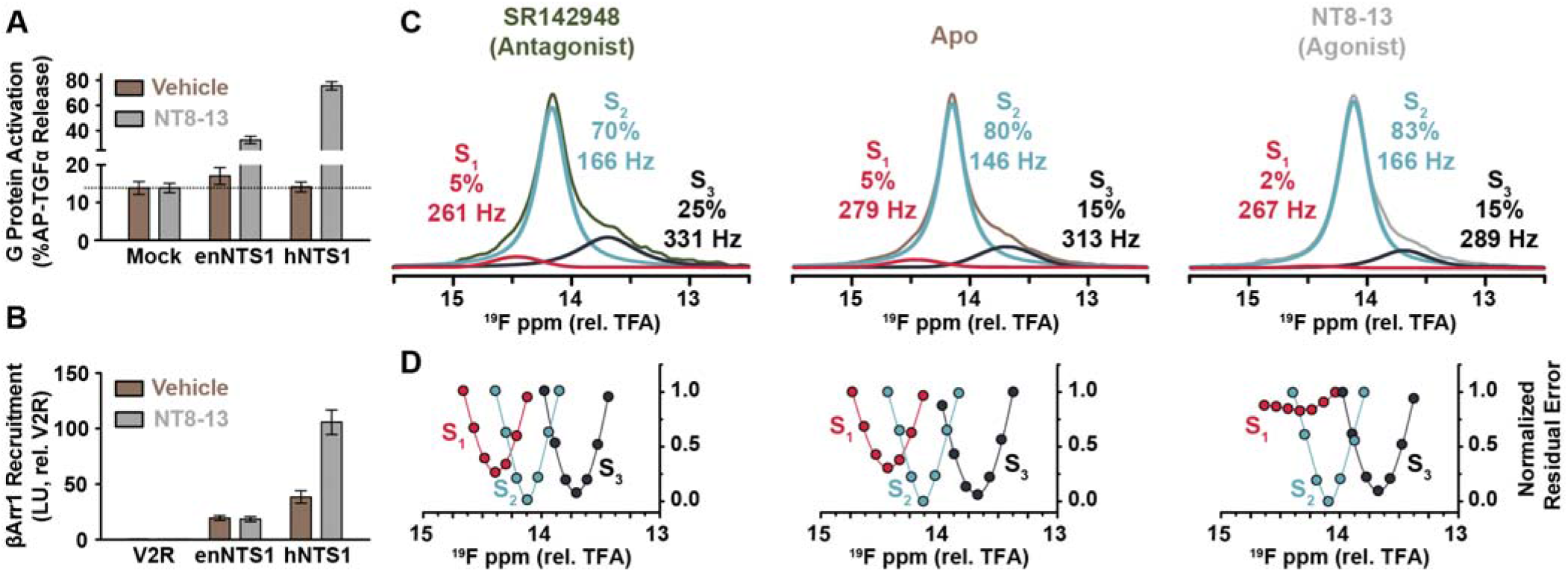
Orthosteric ligands modulate the enNTS1 conformational ensemble. (A) G protein activation was assessed using a TGFα shedding assay on HEK293A cells transiently-transfected with vasopressin receptor 2 (V2R; Mock), human (h)NTS1, or enNTS1.^22^ Cells were stimulated with vehicle (brown) or 1 μM NT8-13 (grey). Error bars represent SEM from three independent experiments. (B) βArr1 recruitment to V2R (Mock), hNTS1, and enNTS1 was measured using a NanoBiT-based assay.^23^ Cells were stimulated with vehicle (brown) or 1 μM NT8-13 (grey). Luminescence counts recorded from 5-10 min following stimulation were averaged and normalized to the initial counts. Error bars represent SEM from four independent experiments. (C) Deconvoluted ^19^F-NMR spectra of enNTS1[Q301C^BTFMA^] in various liganded states. All ligands added to receptor at 10 Meq. The relative population and LWHH are indicated for each substate. (D) The chemical shift value of each deconvoluted resonance was confirmed by monitoring the residual error while constraining peak height and LWHH. The chemical shift was constrained to a new value and the procedure repeated. The lowest residual error value for each substate represents the chemical shift used in deconvolution.^24^

It is unclear which enNTS1 thermostabilizing mutations are responsible for attenuating G protein activation and βArr1 recruitment. We reverted stabilizing mutations adjacent to the connector region (V358F^7.42^) and within the sodium binding site (S113D^2.50^/A362S^7.46^), which are considered critical for activity, but neither backmutation recovered signaling in the TGFα shedding assay (data not shown).^16,39^

### ^19^F-NMR probe does not affect enNTS1 function

To characterize enNTS1’s structural ensemble in solution, we developed protocols to selectively-incorporate cysteine-reactive ^19^F-NMR probes onto TM6. Many previous ^19^F-NMR studies of GPCRs target position 6.27 (Ballesteros-Weinstein nomenclature), but coupling the ^19^F-2-Bromo-N-(4-(trifluoromethyl)phenyl)acetamide (BTFMA) probe at this site reduced enNTS1 expression yields and stability (data not shown).^24,40,41^ MtsslWizard was used to model cysteine-conjugated BTFMA probes at various alternative positions along TM6 of the apo (PDB 6Z66), agonist NT8-13-bound (PDB 4BWB), antagonist SR142948-bound (PDB 6Z4Q), Gα_i_βγ protein ternary (PDB 6OS9), and βArr1 ternary NTS1 complex structures (PDB 6UP7 and 6PWC).^9,10,12,13,34,42^ MtsslWizard rapidly screened 200 randomly-generated BTFMA rotamers and enumerated all conformers that did not clash with the receptor to a tolerance of 3.4 Å. While position 6.27 is unrestricted in antagonist and transducer-bound models, the tight TM5/TM6 packing in the apo and agonist-bound structures sterically-restricted BTFMA to 18 and 110 potential rotamers, respectively, suggesting a mechanism for its observed instability (Figure S2A). In contrast, the neighboring residue Q301C^6.28^ presented completely unhindered mobility in all six structural models (Figure S2A and Table S1). BTFMA-labeling at position 6.28 had no effect on receptor thermostability or yield.

In the final construct, herein enNTS1[Q301C^BTFMA^], solvent exposed C172^3.55^ was mutated to serine to prevent off-site labeling. Site-specific BTFMA labeling was confirmed by LC/MS and NMR with estimated efficiencies of >95% and >80%, respectively (Figure S2B,C and Table S2). enNTS1[Q301C^BTFMA^] showed no appreciable difference in affinity for agonist NT8-13 in saturation binding experiments compared to unlabeled enNTS1, indicative of proper receptor folding (Figure S2D). Dynamic NMR experiments require the sample to be stable throughout multiday data acquisition. To confirm enNTS1[Q301C^BTFMA^] would remain viable during extended periods of data collection, we measured its ability to bind fluorescently-labeled NT8-13 as a function of time. After ten days at 37 °C, 55.0 ± 5.7% apo and 82.1 ± 17.1% agonist-bound enNTS1 [Q301C^BTFMA^] preserved binding competency (Figure S3).

### enNTS1’s conformational ensemble is sensitive to orthosteric ligands

We collected 1D ^19^F-NMR spectra of enNTS1[Q301C^BTFMA^] in the absence and presence of saturating (10:1 Meq) orthosteric ligand concentrations to investigate the conformational ensemble; all spectra possessed S/N ratios ranging from 95.5 to 199.1. Spectral deconvolution of ligand-free enNTS1[Q301C^BTFMA^] best-fit three Lorentzian curves, which qualitatively indicates a three-state equilibrium in slow (ms-s) exchange on the NMR timescale (Figure 1C and Figure S4). The area, chemical shift, and linewidth at half-height (LWHH) for each deconvoluted resonance serve as direct reporters of the relative population, chemical environment, and flexibility of each conformer, respectively.^43^ Following the approach established by Prosser and colleagues, best-fit values were identified by individually constraining a given substate’s chemical shift over a range of frequencies and then globally-fitting the remaining parameters (Figure 1D and Figure S4).^24^ A chemical exchange saturation transfer (^19^F-CEST) experiment was performed on apo enNTS1[Q301C^BTFMA^] to further validate the existence of three substates. The region from −800 Hz to +600 Hz (16.67-11.81 ppm), relative to the S_2_ substate, was scanned in 100 Hz increments with 1 s saturation pulses (Figure 2A). Frequency-dependent changes in peak height confirm the existence of three substates undergoing slow conformational exchange on the NMR timescale (Figure 2A).

**Figure 2.**
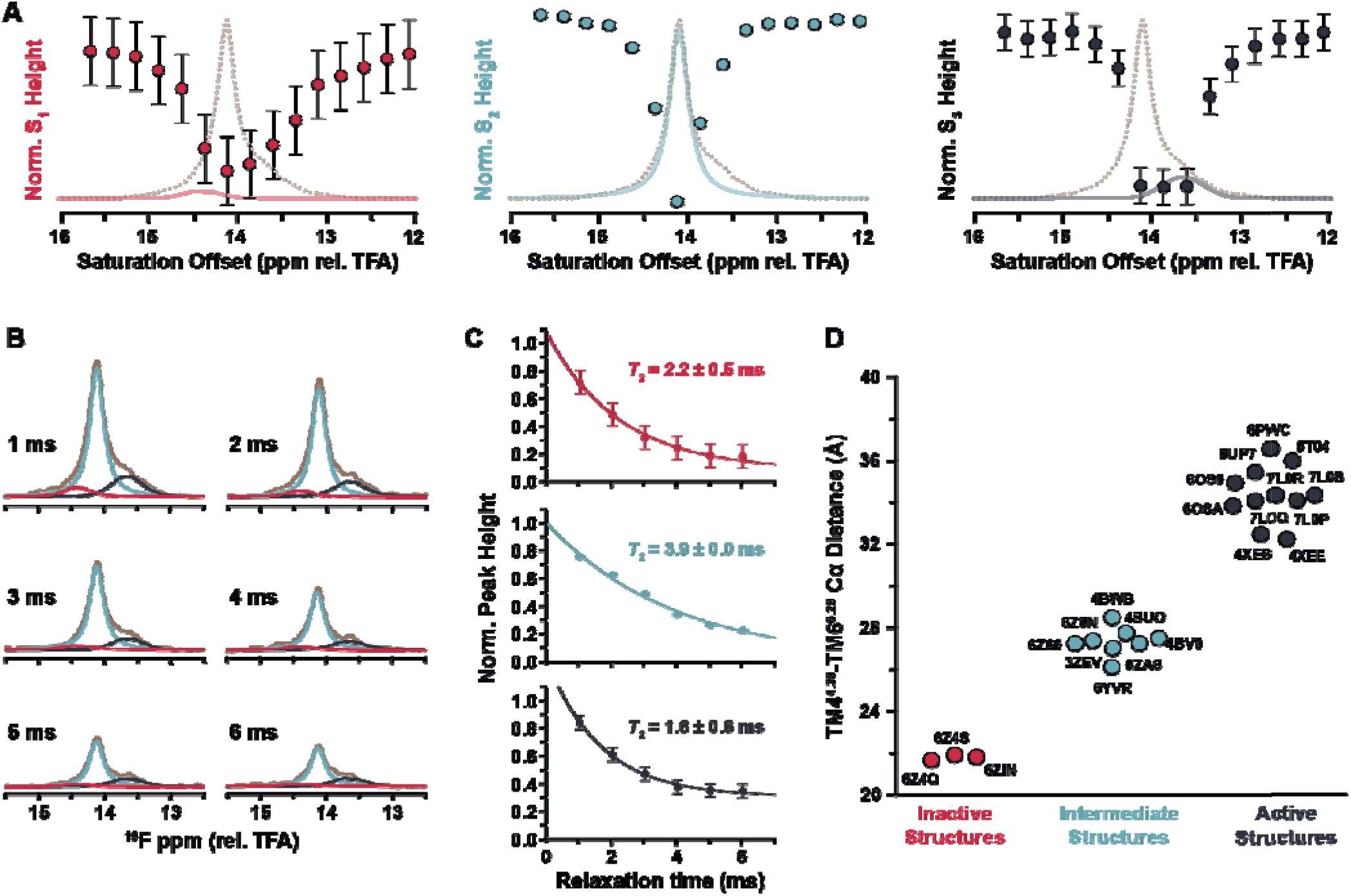
enNTS1 substate exchange and transverse relaxation line broadening. (A) ^19^F-CEST experiments usin**g** apo enNTS1[Q301C^BTFMA^] confirm the presence of three substates qualitatively interconverting on the ms-s timescale. A series of 1D spectra were collected varying the offset frequency of a 1 s saturation pulse in 100 Hz intervals. Spectra were deconvoluted and the height of each substate was normalized to its respective height in the presence of a far off-resonance 1 s control saturation. (B) A series of deconvoluted CPMG T_2_ spectra collected with 1 ms CPMG spin-echo and total relaxation delay varied from 1-6 ms. (C) Fitting the normalize**d** peak heights of the deconvoluted CPMG T_2_ spectra to a mono-exponential model. CEST and CPMG T_2_ error bars represent the standard deviation as calculated at a single offset frequency across the entire spectral series. (D) The TM4-TM6 Cα distance between NTS1 S182^4.38^ and Q301 ^6.28^, plotted for all NTS1 atomic models deposited in the Protein Data Bank, correlates with their putative activation state.

In the apo state, the three resonances (labeled S_1_, S_2_, and S_3_) were populated at 5%, 80%, and 15%, respectively, with LWHH ranging from 146-313 Hz (Figure 1C). For the deconvoluted ^19^F-1D spectra, LWHH = 1/πT_2_*, where T_2_* = T_2,homogenous_ + T_2,inhomogenous_. T_2,homogenous_ is the natural linewidth modulated by μs-ms chemical exchange whereas T_2,inhomogenous_ result from magnetic field inhomogeneities and qualitatively slower chemical exchange processes.^44,45^ A Carr-Purcell-Meiboom-Gill (CPMG) T_2_ experiment is capable of refocusing T_2,inhomogenous_ and thus, to first approximation, reports only the T_2,homogenous_ contribution.^44,45^ Using a train of ~1 ms CPMG spin-echo periods over a 1-6 ms total relaxation delay, all three substates exhibited a mono-exponential decay in peak height (Figure 2B). Both the LWHH and T_2_ were directly fitted for each substate and compared to the linewidths and T_2_ derived from the 1D deconvolution (Figure 2C and Table S3). We report T_2,inhomogenous_ contributions on the order of 51-62% T_2_ for each substate; assuming a homogenous sample preparation in a well-shimmed modern spectrometer, T_2,inhomogenous_ should be negligible.^46^ Thus, our results suggest that conformational exchange on the order of the millisecond CPMG delay is also being partially refocused, although rigorous determination would require relaxation dispersion-type CPMG experiments.^46^

The same three substates were also present in agonist- and antagonist-bound spectra; agonist reduced the S_1_ population while increasing S_2_, whereas antagonist had the opposite effect (Figure 1C and Figure S4). Both ligands similarly decreased the S_1_ LWHH ~20 Hz suggesting a slight stabilizing effect. The S_2_ substate exhibited subtle ligand-dependent frequency perturbations – shifting approximately 0.01 ppm downfield and 0.04 ppm upfield in response to SR142948 antagonist and NT8-13 agonist, respectively (Figure 1C). The simplest explanation for this behavior is that the metastable S_2_ substate is in fast exchange between two high energy microstates, such as local stereoisomers, where the S_2_ chemical shift reflects the relative population of each microstate.^46^ These modest chemical shift perturbations were accompanied by ~20 Hz line broadening. Similarly, the S_3_ linewidth reported on ligand-efficacy with agonist decreasing, and antagonist increasing, the LWHH by 20 Hz (Figure 1C).

### Orthosteric ligands modulate distinct conformational kinetics

The simultaneous observation of three distinct enNTS1[Q301C^BTFMA^] resonances defines an upper limit of approximately 10^-3^ s^-1^ to the exchange rates. We undertook saturation transfer difference (STD) dynamic NMR experiments to quantify the exchange kinetics between substates. ^19^F-STD experiments employ a low power pulse to selectively saturate (i.e. reduce the intensity) a single substate frequency, *v_s_*. When a saturated substate exchanges, it decreases the signal at the other site(s). A series of ^19^F-1D spectra were collected with the saturation pulse duration varied from 50-1000 ms. To account for off-resonance saturation effects, a second series of ^19^F-1D spectra were collected with a control saturation pulse set at an equal, but opposite, offset (*v_c_*) from the substate of interest (Figure S5). The difference in peak height between on- and off-resonance experiments (*v*_*s*,eff_), as a function of saturation pulse length, can be fitted to the Bloch-McConnell equations to yield the exchange rate constant (*k*) with the irradiated resonance, and by extension the lifetimes (*τ_s_* = 1/*k*) of each conformer.^47^ Judicious selection of irradiation frequencies is paramount for minimization of off-resonance artifacts and incomplete saturation, but the limited spectral dispersion of enNTS1[Q301C^BTFMA^] substates presented an insurmountable experimental challenge that underlies ambiguity in the accuracy of fitted exchange rates (Figure S6 and Table **S4**).^48–50^ Nonetheless, we were unable to observe direct exchange between S_1_ and S_3_ under any condition, which supports a linear activation trajectory (S_1_→S_2_→S_3_) from the inactive conformer to the most solvent exposed position (Figure S7, and Table S4). Such a sequential transition was also observed for ^19^F-TM6^6.31^ of the adenosine A_2A_ receptor in LMNG micelles.^51^ Although, subsequent studies in nanodiscs resolved these resonances into two distinct nucleotide-exchange competent states, and an activation intermediate, complicate this comparison with enNTS1 [Q301C^BTFMA^].^52^ As an orthogonal estimate of exchange rates, we collected a 2D [^19^F,^19^F]-exchange spectroscopy (EXSY) spectrum on apo enNTS1[Q301C^BTFMA^] with a 100 ms mixing time. Weak diagonal peaks were only observed for S_2_ and S_3_ substates, but no exchange cross peaks were visible likely due to the overall poor S/N ratio (Figure S8).

Altogether, we hypothesize that S_1_ is an inactive conformation whereas S_2_ and S_3_ reflect active-intermediate and active-like states, respectively. This is based upon several similar observations with other ^19^F-TM6 labeled receptors: i) the comparatively broad S_1_ linewidth, which is consistent with μs-ms timescale motions such as DRY ionic-lock flickering between formation/disruption reported for the β_2_-, β_1_-adrenergic, and A_2A_ adenosine receptors;^53–55^ ii) the near disappearance of the S_1_ resonance and concurrent increase of the S_2_ population upon agonist addition; and iii) the S_1_, S_2_, and S_3_ substate chemical shifts are increasingly up-field, which is consistent with increased solvent exposure as the cytoplasmic cavity expands for transducer association.^41,56^ Although likely an oversimplification, we speculate that the S_1_, S_2_, and S_3_ substates represent the three conformations that all 24 NTS1 atomic models can be organized into based upon the intracellular TM4-TM6 distance (Figure 2D).

### G protein mimetic stabilizes novel conformations

Next, we investigated the interaction of enNTS1 with a synthetic peptide (herein Gα_q_ peptide) corresponding to residues 333-359 of the Gα_q_ C-terminus (a.k.a. α5-helix). The α5-helix is conserved across Gα protein subunits as a random coil that adopts a helical structure upon receptor recognition that comprises 55-69% of the GPCR/G protein interface surface area.^57–59^ We first characterized the efficacy of the enNTS1/Gα_q_ peptide interaction using an affinity pulldown approach.^30^ The Gα_q_ peptide was N-terminally fused to a biotin tag and enNTS1 contained a C-terminal monomeric, ultra-stabilized green fluorescent protein (muGFP) fusion (enNTS1-muGFP).^60^ Binding efficacy was quantified as the fluorescence ratio of streptavidin-captured enNTS1-muGFP versus total (i.e. streptavidin-captured plus unbound) fluorescence.

In the absence of a ligand, the Gα_q_ peptide captured 26.2 ± 6.4% apo enNTS1-muGFP (Figure 3A). Repeating the pulldown in the presence of saturating NT8-13 agonist increased the enNTS1-muGFP capture efficiency to 36.4 ± 6.6% whereas the SR142948 antagonist had no significant effect. Performing the experiment with enNTS1[Q301C^BTFMA^] showed no appreciable differences from enNTS1, indicating that the TM6 ^19^F-BTFMA label does not influence Gα_q_ peptide interaction (Figure 3A). Assuming a quadratic binding model, our results suggest an enNTS1[Q301C^BTFMA^]/Gα_q_ peptide complex K_d_ ~ 225 μM, or a higher basal-state association as reflected in the TGFα shedding assay (Figure 1A).

**Figure 3.**
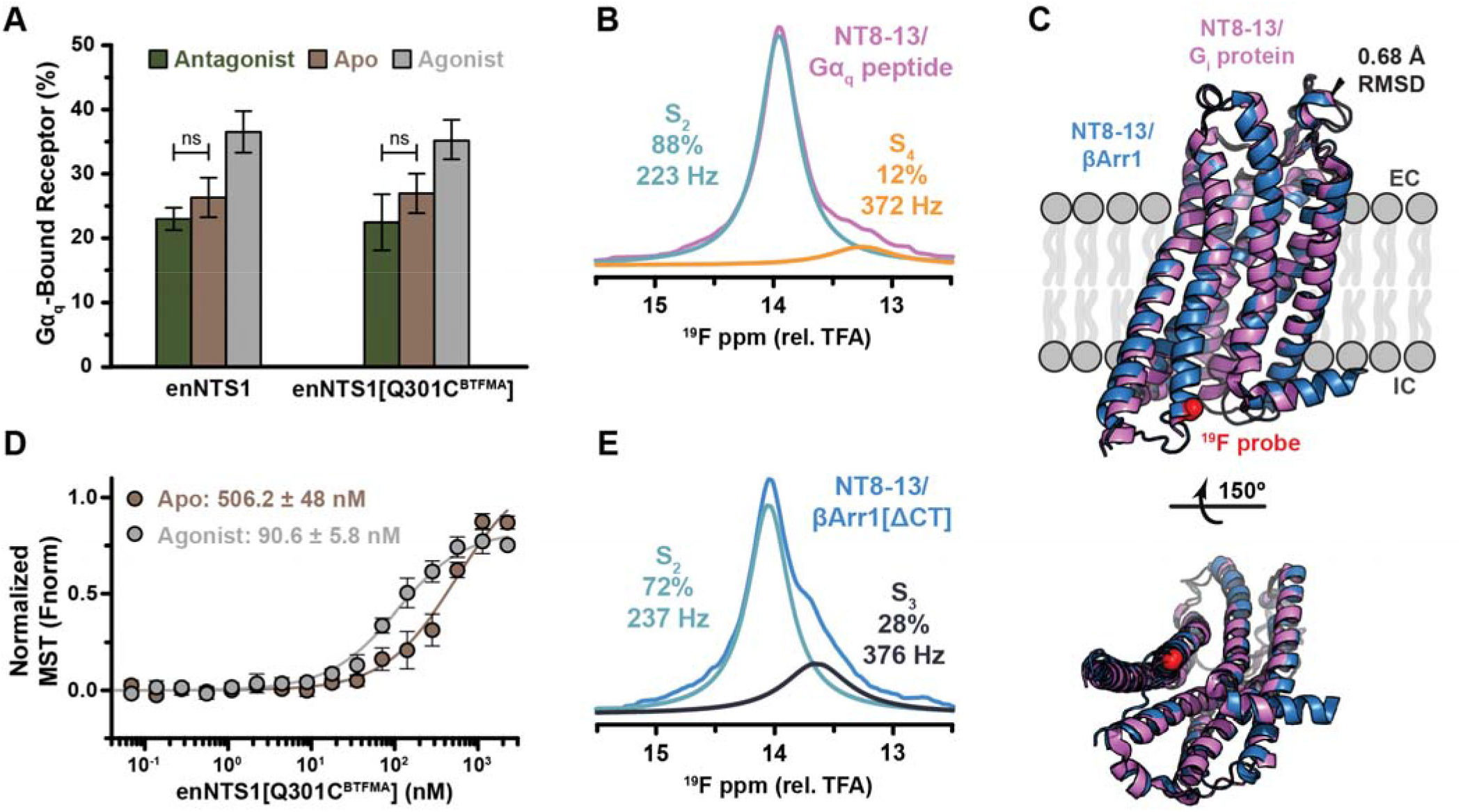
Gα_q_ peptide and βArr1[ΔCT] stabilize distinct enNTS1 substates. (A) Binding of the Gα_q_ peptide to enNTS1 and enNTS1[Q301C^BTFMA^] was measured using a Dynabead sequestration assay.^30^ Ligands were incubated at 10 Meq and Gα_q_-bound receptor calculated as the ratio of input/bound receptor. Bars represent the average bound percentage from both experimental and instrumental triplicates; error bars represent the standard deviation. Statistical significance between conditions was calculated at P = 0.05 using a one-way ANOVA test; “ns” denotes no significance between two conditions as determined via calculated F-ratio at P = 0.05. (B) Deconvoluted ^19^F-NMR spectra of enNTS1[Q301C^BTFMA^] in the presence of 10 Meq NT8-13 and 5 Meq G_αq_ peptide. Two Lorentzian lineshapes (S_2_ and S_4_) provided the best fit to the experimental data. (C) Overlay of NTS1 receptor from NT8-13/heterotrimeric G_i_ protein (PDB 6OS9; magenta) and NT8-13/βArr1 (PDB 6UP7; blue) complex structures. Transducer and agonist were removed for clarity. The distance between Q301 in the two structures is 0.8 Å and the all-atom RMSD = 0.68 Å as calculated using PYMOL. (D) The affinity (K_d_) of NT8-13-bound enNTS1[Q301C^BTFMA^] for βArr1[ΔCT] was measured using microscale thermophoresis (MST). Fluorescent NTA-labeled βArr1[ΔCT] (25 nM), NT8-13 (22.5 μM), and PIP_2_ (10 Meq) were incubated with increasing concentrations of enNTS1[Q301C^BTFMA^]. Data points represent the average normalized MST signal from data collected in both experimental and instrumental triplicates; error bars represent the standard deviation. Equilibrium dissociation constants were calculated from a global fit of experimental data using the quadratic binding model. (E) Deconvoluted ^19^F-NMR spectra of enNTS1[Q301C^BTFMA^] in the presence of 10 Meq NT8-13, 10 Meq PIP_2_, and 5 Meq βArr1 [ΔCT]. Two Lorentzian lineshapes (S_2_ and S_3_) provided the best fit to the experimental data.

The Gα_q_ peptide modified the NT8-13-bound enNTS1[Q301C^BTFMA^] ^19^F-NMR spectra by replacing the inactive S_1_ substate, with a unique peak at 13.26 ppm (S_4_) alongside S_2_ (Figure 3B, Figure S9, and Figure S10). Relative to NT8-13 alone, formation of the Gα_q_ peptide ternary complex increases the S_2_ population from 84% to 88%, broadens the linewidth 57 Hz, and perturbs the chemical shift an additional 0.11 ppm upfield. If we assume the frequency difference between the two pure S_2_ microstates ⍰ 100 Hz (0.18 ppm), the exchange process is likely on the low millisecond timescale although relaxation dispersion-type CPMG experiments are required for quantification.^46^ The Gα_q_ peptide-induced S_4_ substate is upfield of any ligand-only conformer consistent with previous studies that agonist alone is unable to completely stabilize the fully active conformation.^21^ It is possible that the S_4_ substate exists as a broad underlying resonance that is undetectable in the absence of Gα_q_ peptide. If so, we would anticipate observation in the CEST experiment although S/N could be limiting (Figure 2A). The S_4_ resonance linewidth is 372 Hz and constitutes 12% of the observed populations (Figure 3B,E). It is also possible that the broad linewidth is a result of multiple overlapping resonances, but there is insufficient evidence to deconvolute additional conformers.

A similar distribution of TM6 G protein-bound conformers has also been observed for the adenosine A_2A_ receptor in complex with Gα_s_ peptide via ^19^F-NMR.^51^ Once bound to the stimulatory G protein peptide and cognate agonist, TM6 populated two distinct conformers at the expense of all inactive substates. A concurrent population increase for the upfield-most chemical shift occurred, similar to S_4_ for enNTS1[Q301C^BTFMA^]. It is possible that for a majority of class-A GPCRs, complexation with G proteins induce μs-ms timescale chemical exchange of TM6 reflecting pre-coupling conformations prior to full receptor stimulation.

### Arrestin stabilizes pre-existing conformations

Recent cryo-EM structures of hNTS1/βArr1 and hNTS1/G_i_ protein reveal a remarkably conserved receptor architecture with a 0.67 Å all-atom RMSD (Figure 3C).^10,12^ We next wanted to test if βArr1 modified the enNTS1 intracellular landscape similar to the Gα_q_ peptide. βArr1 recruitment is physiologically-dependent on receptor phosphorylation, primarily on intracellular loop 3 (ICL3) and the C-terminus, but the number and location of sites necessary and sufficient to promote coupling is relatively unknown.^61^ To reduce system complexity and facilitate a high-affinity interaction, we employed a preactivated human βArr1 variant truncated at N382 (herein βArr1[ΔCT]).^31^ Microscale thermophoresis (MST) was used to determine the apparent equilibrium dissociation constants (K_d_) of enNTS1[Q301C^BTFMA^]/βArr1[ΔCT] complexes. The N-terminal His-tag of βArr1[ΔCT] was site-specifically labeled with the RED-tris-NTA 2^nd^ Generation (Monolith) fluorescent dye. RED-βArr1[ΔCT] was then incubated with increasing enNTS1[Q301C^BTFMA^] concentrations in the presence or absence of saturating NT8-13 agonist. The interactions followed a sigmoidal dose-response and affinities were calculated using the quadratic binding model. Apo enNTS1[Q301C^BTFMA^] bound RED-βArr1[ΔCT] with a K_d_ ≥ 506.2 ± 48 nM consistent with the high-affinity reported for pre-activated arrestin variants (Figure 3D).^10,62^ The NT8-13 agonist increased affinity to 90.6 ± 5.8 nM, which is similar to the NTS1/Gα_i_βγ ternary complex in phospholipid nanodiscs.^14^

Analogous to the Gα_q_ peptide, βArr1[ΔCT] abolished the inactive S_1_ substate (Figure 3E, Figure S9, and Figure S10). But rather than inducing a new substate resonance, βArr1[ΔCT] selectively restructured the existing conformational landscape by increasing the S_3_ population to 28% and decreasing S_2_ to 72% (Figure 3D). The linewidths of S_2_ and S_3_ increased to 237 Hz and 376 Hz, respectively, suggesting additional contributions of μs-ms timescale chemical exchange (Figure 3E).

## CONCLUSION

Provided that the distance between the TM4 and TM6 intracellular tips approximates transducer binding competency, Figure 2D illustrates that all NTS1 structures determined to date can be organized into three functional categories (inactive, active-intermediate, and active). Our spectroscopic results demonstrate that enNTS1 dynamically populates an ensemble of at least four conformers that are allosterically-tuned by the orthosteric pocket. It is likely that some of these ^19^F-TM6 substates correspond directly to the static structures; future experiments could validate interhelical distances using electron paramagnetic resonance (EPR) or florescence spectroscopy.

As shown for other Class-A GPCRs, agonist binding does not stabilize a single enNTS1 active-like conformation but rather tunes the energetic landscape (Figure 4).^21^ Congruently, βArr1[ΔCT] selects from pre-existing active-like NTS1 substates (Figure 3D). As observed in published atomic models of NTS1/βArr1, steric clash between TM6 and the βArr1 finger-loop eliminates the inactive conformer.^10,13^ The Gα_q_ peptide also reduces the inactive substate while inducing one active-like substate distinct from those observed in the presence of orthosteric ligands or βArr1 (Figure 3B). The Gα_i_ α5-helix peptide had a similar effect on the conformational ensemble of a related thermostabilized NTS1 variant.^63^ Both transducer mimetics increased the active-intermediate and active-like linewidths by 130% relative to NT8-13. This line broadening likely reflects additional dynamics associated with encounter complex formation and the established inherent dynamics of both molecules.^14,52,64–66^

**Figure 4.**
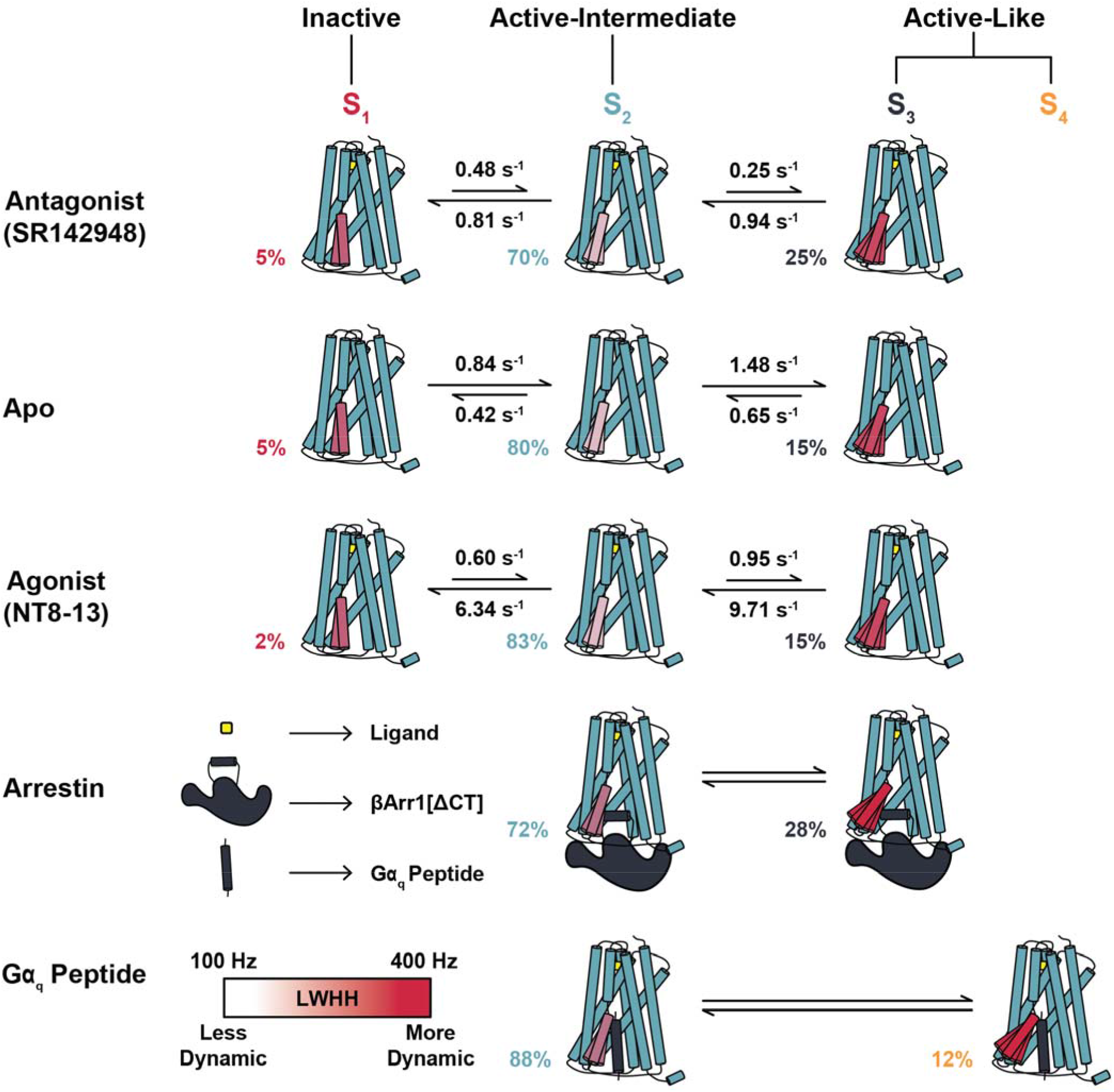
Model summarizing the effect of orthosteric ligands and transducers on each enNTS1 substate lifetime, population, and dynamics.

While both transducers dock helical motifs into the receptor’s cytosolic core, the orientation of each segment is structurally unique; the inserted βArr1 finger-loop is 90° relative to TM6 whereas the G protein α5-helix is parallel.^10,12^ These subtly distinct interfaces may promote structural fluctuations of TM6 that are not easily captured in static models. Together, this suggests that the enNTS1 allosteric activation mechanism may alternate between induced fit (Gα_q_) and conformational selection (βArr1) depending on the coupled transducer. Employing a nucleotide-free heterotrimeric G protein and a phosphorylation-mediated βArr1 ternary complex, which represent “end point” conformations, should enable path delineation. It is important to note the inherent challenge of extrapolating observations at a single probe site to global conformational dynamics. Yet, the sensitivity of ^19^F-NMR to distinct ternary complexes, despite highly similar architecture, suggests that varying the probe location around the helical bundle will provide unprecedent insight of the lowly-populated states of the NTS1 ensemble.^10,12^

An ensemble view in which transducers and ligands modulate the thermodynamic populations and exchange kinetics of enNTS1 substates provides a foundation for designing molecules that select discrete transducer pathways. The relatively recent recognition of this so called biased signaling, in which ligands preferentially-activate either the G protein or arrestin pathway, offers a new mechanism for reducing drug side effects.^67–69^ There are several promising biased agents in pre-clinical and clinical trials; most notably the opioid receptor G protein-biased ligand oliceridine (TRV130) which was approved for pain management in August 2020.^70,71^ A class of βArr1-biased agents have also been developed that target NTS1 and attenuate methamphetamine and cocaine abuse while limiting G protein-mediated on-target side effects.^72^ Our results suggest that ^19^F-NMR may serve as a powerful discovery platform to delineate biased agonists and biased allosteric modulators.

An important limitation of our model system is the employment of a thermostabilized receptor and pre-activated transducer mimetics. Although commonly employed to stabilize a single conformational state for structure determination, these mutations rigidify receptor motions and alter specific inter-residue and receptor/solvent interactions.^73,74^ Nonetheless, enNTS1 responds appropriately, although with reduced efficacy, to ligands and allosteric modulators in functional assays. Given that mutations are more commonly loss-of-function rather than gain-of-function, we hypothesize that thermostabilization does not result in an entirely distinct activation landscape. While more experiments are required to further the dynamic NTS1 model presented here, this study illustrates the importance of orthogonal structural techniques in understanding the complete mechanism of GPCR activation.

## ASSOCIATED CONTENT

### Supporting Information

The Supporting Information is available free of charge on the ACS Publications website.

Detailed experimental procedures (PDF)

## AUTHOR INFORMATION

### Notes

The authors declare no competing financial interests.

## ACKNOWLEDGMENT

We are grateful to Prof. Gregor Hagelueken at the University of Bonn for assistance and modeling in the mtsslWizard software, Dr. Hongwei Wu at Indiana University for NMR instrument assistance, Dr. Ratan Rai at Indiana University School of Medicine for NMR instrument assistance, Kouki Kawakami at Tohoku University for technical assistance, Kayo Sato, Shigeko Nakano and Ayumi Inoue at Tohoku University for their assistance in plasmid preparation and cell-based GPCR assays, Prof. Ashish Manglik at the University of California for providing the βArr1 construct used in this study, and Prof. Daniel Scott at the Florey Institute for providing the enNTS1 plasmid used in this study. The 14.1 T spectrometers used in this study were generously supported by the Indiana University Fund.

---

Insert Table of Contents artwork here

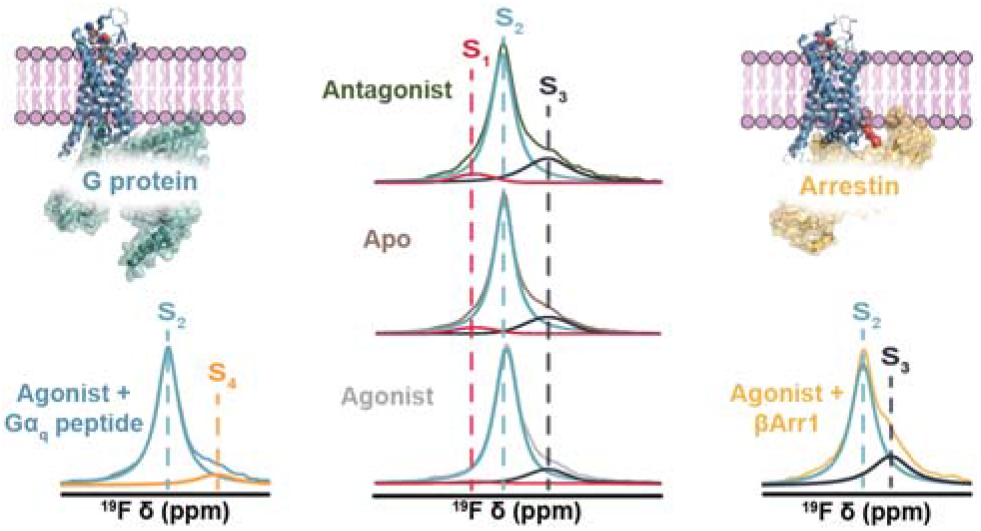

## REFERENCE

(1) de Mendoza, A.; Sebé-Pedrós, A.; Ruiz-Trillo, I. The Evolution of the GPCR Signaling System in Eukaryotes: Modularity, Conservation, and the Transition to Metazoan Multicellularity. Genome Biol. Evol. 2014, 6 (3), 606–619. https://doi.org/10.1093/gbe/evu038.

(2) Hanlon, C. D.; Andrew, D. J. Outside-in Signaling – a Brief Review of GPCR Signaling with a Focus on the Drosophila GPCR Family. J. Cell Sci. 2015, 128 (19), 3533–3542. https://doi.org/10.1242/jcs.175158.

(3) Hauser, A. S.; Attwood, M. M.; Rask-Andersen, M.; Schiöth, H. B.; Gloriam, D. E. Trends in GPCR Drug Discovery: New Agents, Targets and Indications. Nat. Rev. Drug Discov. 2017, 16, 829.

(4) Munk, C.; Isberg, V.; Mordalski, S.; Harpsøe, K.; Rataj, K.; Hauser, A. S.; Kolb, P.; Bojarski, A. J.; Vriend, G.; Gloriam, D. E. GPCRdb: The G Protein-Coupled Receptor Database - an Introduction. Br. J. Pharmacol. 2016, 173 (14), 2195–2207. https://doi.org/10.1111/bph.13509.

(5) Grisshammer, R. New Approaches towards the Understanding of Integral Membrane Proteins: A Structural Perspective on G Protein-Coupled Receptors. Protein Sci. 2017, 26 (8), 1493–1504. https://doi.org/10.1002/pro.3200.

(6) Schönegge, A.-M.; Gallion, J.; Picard, L.-P.; Wilkins, A. D.; Le Gouill, C.; Audet, M.; Stallaert, W.; Lohse, M. J.; Kimmel, M.; Lichtarge, O.; Bouvier, M. Evolutionary Action and Structural Basis of the Allosteric Switch Controlling B2AR Functional Selectivity. Nat. Commun. 2017, 8 (1), 2169. https://doi.org/10.1038/s41467-017-02257-x.

(7) Sanchez-Soto, M.; Verma, R. K.; Willette, B. K. A.; Gonye, E. C.; Moore, A. M.; Moritz, A. E.; Boateng, C. A.; Yano, H.; Free, R. B.; Shi, L.; Sibley, D. R. A Structural Basis for How Ligand Binding Site Changes Can Allosterically Regulate GPCR Signaling and Engender Functional Selectivity. Sci. Signal. 2020, 13 (617), aaw5885. https://doi.org/10.1126/scisignal.aaw5885.

(8) Zhou, Q.; Yang, D.; Wu, M.; Guo, Y.; Guo, W.; Zhong, L.; Cai, X.; Dai, A.; Jang, W.; Shakhnovich, E. I.; Liu, Z.-J.; Stevens, R. C.; Lambert, N. A.; Babu, M. M.; Wang, M.-W.; Zhao, S. Common Activation Mechanism of Class A GPCRs. Elife 2019, 8, e50279–e50279. https://doi.org/10.7554/eLife.50279.

(9) Deluigi, M.; Klipp, A.; Klenk, C.; Merklinger, L.; Eberle, S. A.; Morstein, L.; Heine, P.; Peer, M. R. E.; Ernst, P.; Kamenecka, T. M.; He, Y.; Vacca, S.; Egloff, P.; Honegger, A.; Pluckthun, A. Complexes of the Neurotensin Receptor 1 with Small-Molecule Ligands Reveal Structural Determinants of Full, Partial, and Inverse Agonism. Sci. Adv. 2021, 7 (5), eabe5504–eabe5504. https://doi.org/10.1126/sciadv.abe5504.

(10) Huang, W.; Masureel, M.; Qu, Q.; Janetzko, J.; Inoue, A.; Kato, H. E.; Robertson, M. J.; Nguyen, K. C.; Glenn, J. S.; Skiniotis, G.; Kobilka, B. K. Structure of the Neurotensin Receptor 1 in Complex with β-Arrestin 1. Nature 2020, 579 (7798), 303–308. https://doi.org/10.1038/s41586-020-1953-1.

(11) Egloff, P.; Hillenbrand, M.; Klenk, C.; Batyuk, A.; Heine, P.; Balada, S.; Schlinkmann, K. M.; Scott, D. J.; Schütz, M.; Plückthun, A. Structure of Signaling-Competent Neurotensin Receptor 1 Obtained by Directed Evolution in Escherichia Coli. Proc. Natl. Acad. Sci. 2014, 111 (6), E655 LP–E662.

(12) Kato, H. E.; Zhang, Y.; Hu, H.; Suomivuori, C.-M.; Kadji, F. M. N.; Aoki, J.; Krishna Kumar, K.; Fonseca, R.; Hilger, D.; Huang, W.; Latorraca, N. R.; Inoue, A.; Dror, R. O.; Kobilka, B. K.; Skiniotis, G. Conformational Transitions of a Neurotensin Receptor 1–Gi1 Complex. Nature 2019, 572 (7767), 80–85. https://doi.org/10.1038/s41586-019-1337-6.

(13) Yin, W.; Li, Z.; Jin, M.; Yin, Y.-L.; de Waal, P. W.; Pal, K.; Yin, Y.; Gao, X.; He, Y.; Gao, J.; Wang, X.; Zhang, Y.; Zhou, H.; Melcher, K.; Jiang, Y.; Cong, Y.; Edward Zhou, X.; Yu, X.; Eric Xu, H. A Complex Structure of Arrestin-2 Bound to a G Protein-Coupled Receptor. Cell Res. 2019, 29 (12), 971–983. https://doi.org/10.1038/s41422-019-0256-2.

(14) Zhang, M.; Gui, M.; Wang, Z.-F.; Gorgulla, C.; Yu, J. J.; Wu, H.; Sun, Z. J.; Klenk, C.; Merklinger, L.; Morstein, L.; Hagn, F.; Plückthun, A.; Brown, A.; Nasr, M. L.; Wagner, G. Cryo-EM Structure of an Activated GPCR–G Protein Complex in Lipid Nanodiscs. Nat. Struct. Mol. Biol. 2021, 28 (3), 258–267. https://doi.org/10.1038/s41594-020-00554-6.

(15) White, J. F.; Noinaj, N.; Shibata, Y.; Love, J.; Kloss, B.; Xu, F.; Gvozdenovic-Jeremic, J.; Shah, P.; Shiloach, J.; Tate, C. G.; Grisshammer, R. Structure of the Agonist-Bound Neurotensin Receptor. Nature 2012, 490, 508.

(16) Krumm, B. E.; White, J. F.; Shah, P.; Grisshammer, R. Structural Prerequisites for G-Protein Activation by the Neurotensin Receptor. Nat. Commun. 2015, 6, 7895.

(17) Krumm, B. E.; Lee, S.; Bhattacharya, S.; Botos, I.; White, C. F.; Du, H.; Vaidehi, N.; Grisshammer, R. Structure and Dynamics of a Constitutively Active Neurotensin Receptor. Sci. Rep. 2016, 6, 38564. https://doi.org/10.1038/srep38564.

(18) Qiu, S.; Pellino, G.; Fiorentino, F.; Rasheed, S.; Darzi, A.; Tekkis, P.; Kontovounisios, C. A Review of the Role of Neurotensin and Its Receptors in Colorectal Cancer. Gastroenterol. Res. Pract. 2017, 2017, 6456257. https://doi.org/10.1155/2017/6456257.

(19) Remaury, A.; Vita, N.; Gendreau, S.; Jung, M.; Arnone, M.; Poncelet, M.; Culouscou, J.-M.; Le Fur, G.; Soubrié, P.; Caput, D.; Shire, D.; Kopf, M.; Ferrara, P. Targeted Inactivation of the Neurotensin Type 1 Receptor Reveals Its Role in Body Temperature Control and Feeding Behavior but Not in Analgesia. Brain Res. 2002, 953 (1), 63–72. https://doi.org/https://doi.org/10.1016/S0006-8993(02)03271-7.

(20) Boules, M.; Li, Z.; Smith, K.; Fredrickson, P.; Richelson, E. Diverse Roles of Neurotensin Agonists in the Central Nervous System. Frontiers in Endocrinology. 2013, p 36.

(21) Weis, W. I.; Kobilka, B. K. The Molecular Basis of G Protein-Coupled Receptor Activation. Annu. Rev. Biochem. 2018, 87, 897–919. https://doi.org/10.1146/annurev-biochem-060614-033910.

(22) Inoue, A.; Ishiguro, J.; Kitamura, H.; Arima, N.; Okutani, M.; Shuto, A.; Higashiyama, S.; Ohwada, T.; Arai, H.; Makide, K.; Aoki, J. TGFα Shedding Assay: An Accurate and Versatile Method for Detecting GPCR Activation. Nat. Methods 2012, 9 (10), 1021–1029. https://doi.org/10.1038/nmeth.2172.

(23) Dixon, A. S.; Schwinn, M. K.; Hall, M. P.; Zimmerman, K.; Otto, P.; Lubben, T. H.; Butler, B. L.; Binkowski, B. F.; Machleidt, T.; Kirkland, T. A.; Wood, M. G.; Eggers, C. T.; Encell, L. P.; Wood, K. V. NanoLuc Complementation Reporter Optimized for Accurate Measurement of Protein Interactions in Cells. ACS Chem. Biol. 2016, 11 (2), 400–408. https://doi.org/10.1021/acschembio.5b00753.

(24) Kim, T. H.; Chung, K. Y.; Manglik, A.; Hansen, A. L.; Dror, R. O.; Mildorf, T. J.; Shaw, D. E.; Kobilka, B. K.; Prosser, R. S. The Role of Ligands on the Equilibria Between Functional States of a G Protein-Coupled Receptor. J. Am. Chem. Soc. 2013, 135 (25), 9465–9474. https://doi.org/10.1021/ja404305k.

(25) Chen, H.; Viel, S.; Ziarelli, F.; Peng, L. 19F NMR: A Valuable Tool for Studying Biological Events. Chem. Soc. Rev. 2013, 42 (20), 7971–7982. https://doi.org/10.1039/C3CS60129C.

(26) Kitevski-LeBlanc, J. L.; Evanics, F.; Prosser, R. S. Approaches for the Measurement of Solvent Exposure in Proteins by 19F NMR. J. Biomol. NMR 2009, 45 (3), 255. https://doi.org/10.1007/s10858-009-9359-2.

(27) Chao, F.-A.; Byrd, R. A. Protein Dynamics Revealed by NMR Relaxation Methods. Emerg. Top. life Sci. 2020, 2 (1), 93–105. https://doi.org/10.1042/etls20170139.

(28) Göbl, C.; Madl, T.; Simon, B.; Sattler, M. NMR Approaches for Structural Analysis of Multidomain Proteins and Complexes in Solution. Prog. Nucl. Magn. Reson. Spectrosc. 2014, 80, 26–63. https://doi.org/https://doi.org/10.1016/j.pnmrs.2014.05.003.

(29) Kleckner, I. R.; Foster, M. P. An Introduction to NMR-Based Approaches for Measuring Protein Dynamics. Biochim. Biophys. Acta 2011, 1814 (8), 942–968. https://doi.org/10.1016/j.bbapap.2010.10.012.

(30) Gupte, T. M.; Ritt, M.; Dysthe, M.; Malik, R. U.; Sivaramakrishnan, S. Minute-Scale Persistence of a GPCR Conformation State Triggered by Non-Cognate G Protein Interactions Primes Signaling. Nat. Commun. 2019, 10 (1), 4836. https://doi.org/10.1038/s41467-019-12755-9.

(31) Vishnivetskiy, S. A.; Schubert, C.; Climaco, G. C.; Gurevich, Y. V; Velez, M. G.; Gurevich, V. V. An Additional Phosphate-Binding Element in Arrestin Molecule. Implications for the Mechanism of Arrestin Activation. J. Biol. Chem. 2000, 275 (52), 41049–41057. https://doi.org/10.1074/jbc.M007159200.

(32) Bumbak, F.; Keen, A. C.; Gunn, N. J.; Gooley, P. R.; Bathgate, R. A. D.; Scott, D. J. Optimization and 13CH3 Methionine Labeling of a Signaling Competent Neurotensin Receptor 1 Variant for NMR Studies. Biochim. Biophys. Acta - Biomembr. 2018, 1860 (6), 1372–1383. https://doi.org/https://doi.org/10.1016/j.bbamem.2018.03.020.

(33) White, J. F.; Grisshammer, R. Stability of the Neurotensin Receptor NTS1 Free in Detergent Solution and Immobilized to Affinity Resin. PLoS One 2010, 5 (9), e12579–e12579.

(34) Egloff, P.; Hillenbrand, M.; Klenk, C.; Batyuk, A.; Heine, P.; Balada, S.; Schlinkmann, K. M.; Scott, D. J.; Schütz, M.; Plückthun, A. Structure of Signaling-Competent Neurotensin Receptor 1 Obtained by Directed Evolution in Escherichia Coli; Proc. Natl. Acad. Sci. 2014, 111 (6), E655 LP–E662. https://doi.org/10.1073/pnas.1317903111.

(35) White, J. F.; Noinaj, N.; Shibata, Y.; Love, J.; Kloss, B.; Xu, F.; Gvozdenovic-Jeremic, J.; Shah, P.; Shiloach, J.; Tate, C. G.; Grisshammer, R. Structure of the Agonist-Bound Neurotensin Receptor. Nature 2012, 490 (7421), 508–513. https://doi.org/10.1038/nature11558.

(36) García-Garayoa, E.; Bläuenstein, P.; Bruehlmeier, M.; Blanc, A.; Iterbeke, K.; Conrath, P.; Tourwé, D.; Schubiger, P. A. Preclinical Evaluation of a New, Stabilized Neurotensin(8--13) Pseudopeptide Radiolabeled with (99m)Tc. J. Nucl. Med. 2002, 43 (3), 374–383.

(37) Pinkerton, A. B.; Peddibhotla, S.; Yamamoto, F.; Slosky, L. M.; Bai, Y.; Maloney, P.; Hershberger, P.; Hedrick, M. P.; Falter, B.; Ardecky, R. J.; Smith, L. H.; Chung, T. D. Y.; Jackson, M. R.; Caron, M. G.; Barak, L. S. Discovery of β-Arrestin Biased, Orally Bioavailable, and CNS Penetrant Neurotensin Receptor 1 (NTR1) Allosteric Modulators. J. Med. Chem. 2019, 62 (17), 8357–8363. https://doi.org/10.1021/acs.jmedchem.9b00340.

(38) Slosky, L. M.; Bai, Y.; Toth, K.; Ray, C.; Rochelle, L. K.; Badea, A.; Chandrasekhar, R.; Pogorelov, V. M.; Abraham, D. M.; Atluri, N.; Peddibhotla, S.; Hedrick, M. P.; Hershberger, P.; Maloney, P.; Yuan, H.; Li, Z.; Wetsel, W. C.; Pinkerton, A. B.; Barak, L. S.; Caron, M. G. β-Arrestin-Biased Allosteric Modulator of NTSR1 Selectively Attenuates Addictive Behaviors. Cell 2020, 181 (6), 1364–1379.e14. https://doi.org/10.1016/j.cell.2020.04.053.

(39) Dunn, S. M. J.; Davies, M.; Muntoni, A. L.; Lambert, J. J. Mutagenesis of the Rat A1 Subunit of the γ-Aminobutyric Acid Receptor Reveals the Importance of Residue 101 in Determining the Allosteric Effects of Benzodiazepine Site Ligands. Mol. Pharmacol. 1999, 56 (4), 768 LP–774.

(40) Ballesteros, J. A.; Weinstein, H. [19] Integrated Methods for the Construction of Three-Dimensional Models and Computational Probing of Structure-Function Relations in G Protein-Coupled Receptors. In Receptor Molecular Biology; Sealfon, S. C. B. T.-M. in N., Ed.; Academic Press, 1995; Vol. 25, pp 366–428. https://doi.org/https://doi.org/10.1016/S1043-9471(05)80049-7.

(41) Manglik, A.; Kim, T. H.; Masureel, M.; Altenbach, C.; Yang, Z.; Hilger, D.; Lerch, M. T.; Kobilka, T. S.; Thian, F. S.; Hubbell, W. L.; Prosser, R. S.; Kobilka, B. K. Structural Insights into the Dynamic Process of B2-Adrenergic Receptor Signaling. Cell 2015, 161 (5), 1101–1111. https://doi.org/10.1016/j.cell.2015.04.043.

(42) Hagelueken, G.; Ward, R.; Naismith, J. H.; Schiemann, O. MtsslWizard: In Silico Spin-Labeling and Generation of Distance Distributions in PyMOL. Appl. Magn. Reson. 2012, 42 (3), 377–391. https://doi.org/10.1007/s00723-012-0314-0.

(43) Cavanagh, J.; Fairbrother, W. J.; Palmer, A. G.; Rance, M.; Skelton, N. J. Chapter 1 - Classical NMR Spectroscopy; (Second E., Eds.; Academic Press: Burlington, 2007; pp 1–28. https://doi.org/https://doi.org/10.1016/B978-012164491-8/50003-8.

(44) Szyperski, T. Protein NMR Spectroscopy. Encyclopedia of Molecular Cell Biology and Molecular Medicine. 2006. https://doi.org/10.1002/3527600906.mcb.200500055.

(45) Hatada, K.; Kitayama, T. Introduction to NMR Spectroscopy. NMR Spectroscopy of Polymers. 2004, pp 1–42. https://doi.org/10.1007/978-3-662-08982-8_1.

(46) Palmer, A. G. 3rd; Kroenke, C. D.; Loria, J. P. Nuclear Magnetic Resonance Methods for Quantifying Microsecond-to-Millisecond Motions in Biological Macromolecules. Methods Enzymol. 2001, 339, 204–238. https://doi.org/10.1016/s0076-6879(01)39315-1.

(47) McConnell, H. M. Reaction Rates by Nuclear Magnetic Resonance. J. Chem. Phys. 1958, 28 (3), 430–431. https://doi.org/10.1063/1.1744152.

(48) Forsén, S.; Hoffman, R. A. Study of Moderately Rapid Chemical Exchange Reactions by Means of Nuclear Magnetic Double Resonance. J. Chem. Phys. 1963, 39 (11), 2892–2901. https://doi.org/10.1063/1.1734121.

(49) Kingsley, P. B.; Monahan, W. G. Correcting for Incomplete Saturation and Off-Resonance Effects in Multiple-Site Saturation-Transfer Kinetic Measurements. J. Magn. Reson. 2000, 1461, 100–109.

(50) Horst, R.; Horwich, A. L.; Wüthrich, K. Translational Diffusion of Macromolecular Assemblies Measured Using Transverse-Relaxation-Optimized Pulsed Field Gradient NMR. J. Am. Chem. Soc. 2011, 133 (41), 16354–16357. https://doi.org/10.1021/ja206531c.

(51) Ye, L.; Van Eps, N.; Zimmer, M.; Ernst, O. P.; Scott Prosser, R. Activation of the A2A Adenosine G-Protein-Coupled Receptor by Conformational Selection. Nature 2016, 533, 265.

(52) Huang, S. K.; Pandey, A.; Tran, D. P.; Villanueva, N. L.; Kitao, A.; Sunahara, R. K.; Sljoka, A.; Prosser, R. S. Delineating the Conformational Landscape of the Adenosine A(2A) Receptor during G Protein Coupling. Cell 2021, 184 (7), 1884–1894.e14. https://doi.org/10.1016/j.cell.2021.02.041.

(53) Liu, J. J.; Horst, R.; Katritch, V.; Stevens, R. C.; Wüthrich, K. Biased Signaling Pathways in B2-Adrenergic Receptor Characterized by 19F-NMR. Science (80). 2012, 335 (6072), 1106 LP–1110. https://doi.org/10.1126/science.1215802.

(54) Frei, J. N.; Broadhurst, R. W.; Bostock, M. J.; Solt, A.; Jones, A. J. Y.; Gabriel, F.; Tandale, A.; Shrestha, B.; Nietlispach, D. Conformational Plasticity of Ligand-Bound and Ternary GPCR Complexes Studied by 19F NMR of the B1-Adrenergic Receptor. Nat. Commun. 2020, 11 (1), 669. https://doi.org/10.1038/s41467-020-14526-3.

(55) Sušac, L.; Eddy, M. T.; Didenko, T.; Stevens, R. C.; Wüthrich, K. A2A Adenosine Receptor Functional States Characterized by 19F-NMR. Proc. Natl. Acad. Sci. 2018, 115 (50), 12733 LP–12738. https://doi.org/10.1073/pnas.1813649115.

(56) Hull, W. E.; Sykes, B. D. Fluorotyrosine Alkaline Phosphatase. Fluorine-19 Nuclear Magnetic Resonance Relaxation Times and Molecular Motion of the Individual Fluorotyrosines. Biochemistry 1974, 13 (17), 3431–3437. https://doi.org/10.1021/bi00714a002.

(57) Mobbs, J. I.; Belousoff, M. J.; Harikumar, K. G.; Piper, S. J.; Xu, X.; Furness, S. G. B.; Venugopal, H.; Christopoulos, A.; Danev, R.; Wootten, D.; Thal, D. M.; Miller, L. J.; Sexton, P. M. Structures of the Human Cholecystokinin 1 (CCK1) Receptor Bound to Gs and Gq Mimetic Proteins Provide Insight into Mechanisms of G Protein Selectivity. PLOS Biol. 2021, 19 (6), e3001295–e3001295.

(58) Xia, R.; Wang, N.; Xu, Z.; Lu, Y.; Song, J.; Zhang, A.; Guo, C.; He, Y. Cryo-EM Structure of the Human Histamine H1 Receptor/Gq Complex. Nat. Commun. 2021, 12 (1), 2086. https://doi.org/10.1038/s41467-021-22427-2.

(59) Kim, K.; Che, T.; Panova, O.; DiBerto, J. F.; Lyu, J.; Krumm, B. E.; Wacker, D.; Robertson, M. J.; Seven, A. B.; Nichols, D. E.; Shoichet, B. K.; Skiniotis, G.; Roth, B. L. Structure of a Hallucinogen-Activated Gq-Coupled 5-HT(2A) Serotonin Receptor. Cell 2020, 182 (6), 1574–1588.e19. https://doi.org/10.1016/j.cell.2020.08.024.

(60) Scott, D. J.; Gunn, N. J.; Yong, K. J.; Wimmer, V. C.; Veldhuis, N. A.; Challis, L. M.; Haidar, M.; Petrou, S.; Bathgate, R. A. D.; Griffin, M. D. W. A Novel Ultra-Stable, Monomeric Green Fluorescent Protein For Direct Volumetric Imaging of Whole Organs Using CLARITY. Sci. Rep. 2018, 8 (1), 667. https://doi.org/10.1038/s41598-017-18045-y.

(61) Sente, A.; Peer, R.; Srivastava, A.; Baidya, M.; Lesk, A. M.; Balaji, S.; Shukla, A. K.; Babu, M. M.; Flock, T. Molecular Mechanism of Modulating Arrestin Conformation by GPCR Phosphorylation. Nat. Struct. Mol. Biol. 2018, 25 (6), 538–545. https://doi.org/10.1038/s41594-018-0071-3.

(62) Chen, Q.; Perry, N. A.; Vishnivetskiy, S. A.; Berndt, S.; Gilbert, N. C.; Zhuo, Y.; Singh, P. K.; Tholen, J.; Ohi, M. D.; Gurevich, E. V; Brautigam, C. A.; Klug, C. S.; Gurevich, V. V; Iverson, T. M. Structural Basis of Arrestin-3 Activation and Signaling. Nat. Commun. 2017, 8 (1), 1427. https://doi.org/10.1038/s41467-017-01218-8.

(63) Goba, I.; Goricanec, D.; Schum, D.; Hillenbrand, M.; Plückthun, A.; Hagn, F. Probing the Conformation States of Neurotensin Receptor 1 Variants by NMR Site-Directed Methyl Labeling. ChemBioChem 2021, 22 (1), 139–146. https://doi.org/https://doi.org/10.1002/cbic.202000541.

(64) Shiraishi, Y.; Natsume, M.; Kofuku, Y.; Imai, S.; Nakata, K.; Mizukoshi, T.; Ueda, T.; Iwaï, H.; Shimada, I. Phosphorylation-Induced Conformation of B2-Adrenoceptor Related to Arrestin Recruitment Revealed by NMR. Nat. Commun. 2018, 9 (1), 194. https://doi.org/10.1038/s41467-017-02632-8.

(65) Shiraishi, Y.; Kofuku, Y.; Ueda, T.; Pandey, S.; Dwivedi-Agnihotri, H.; Shukla, A. K.; Shimada, I. Biphasic Activation of ß-Arrestin 1 upon Interaction with a GPCR Revealed by Methyl-TROSY NMR. Nat. Commun. 2021, 12 (1), 7158. https://doi.org/10.1038/s41467-021-27482-3.

(66) Keehun, K.; Shayla, P.; Fredrik, S.; M., G. T.; Michael, R.; Matthew, D.; Nagarajan, V.; Sivaraj, S. B2-Adrenoceptor Ligand Efficacy Is Tuned by a Two-Stage Interaction with the Gas C Terminus. Proc. Natl. Acad. Sci. 2021, 118 (11), e2017201118. https://doi.org/10.1073/pnas.2017201118.

(67) Pupo, A. S.; Duarte, D. A.; Lima, V.; Teixeira, L. B.; Parreiras-E-Silva, L. T.; Costa-Neto, C. M. Recent Updates on GPCR Biased Agonism. Pharmacol. Res. 2016, 112, 49–57. https://doi.org/10.1016/j.phrs.2016.01.031.

(68) Ikuo, M.; Olga, O.; M., K. G.; D., J. C.; Keqiang, X.; A., M. K. Distinct Profiles of Functional Discrimination among G Proteins Determine the Actions of G Protein–Coupled Receptors. Sci. Signal. 2015, 8 (405), ra123–ra123. https://doi.org/10.1126/scisignal.aab4068.

(69) Costa-Neto, C. M.; Parreiras-e-Silva, L. T.; Bouvier, M. A Pluridimensional View of Biased Agonism. Mol. Pharmacol. 2016, 90 (5), 587 LP–595. https://doi.org/10.1124/mol.116.105940.

(70) U.S. Food and Drug Administration. FDA Approves New Opioid for Intravenous Use in Hospitals, Other Controlled Clinical Settings https://www.fda.gov/news-events/press-announcements/fda-approves-new-opioid-intravenous-use-hospitals-other-controlled-clinical-settings.

(71) U.S. Food and Drug Administration. Drug Trials Snapshots: OLINVYK https://www.fda.gov/drugs/drug-approvals-and-databases/drug-trials-snapshots-olinvyk.

(72) Liu, J. J.; Horst, R.; Katritch, V.; Stevens, R. C.; Wüthrich, K. Biased Signaling Pathways in B2-Adrenergic Receptor Characterized by 19F-NMR. Science 2012, 335 (6072), 1106–1110. https://doi.org/10.1126/science.1215802.

(73) Tate, C. G. A Crystal Clear Solution for Determining G-Protein-Coupled Receptor Structures. Trends Biochem. Sci. 2012, 37 (9), 343–352. https://doi.org/10.1016/j.tibs.2012.06.003.

(74) Vaidehi, N.; Grisshammer, R.; Tate, C. G. How Can Mutations Thermostabilize G-Protein-Coupled Receptors? Trends Pharmacol. Sci. 2016, 37 (1), 37–46. https://doi.org/10.1016/j.tips.2015.09.005.

