## Supplementary Information for "The Effect of Ligands and Transducers on the Neurotensin Receptor 1 (NTS1) Conformational Ensemble"

**Supporting Information**

Table of Contents:

Experimental Information:

I.	Materials and Methods..................................................................................................................S2

SI Figures.................................................................................................................................................S7

Figure S1. G protein activation and βArr1 recruitment cellular assays......................................................S7

Figure S2. Selective ^19^F-labeling of enNTS1 intracellular helix................................................................S8

Figure S3. Long-term stability of enNTS1[Q301C^BTFMA^].........................................................................S9

Figure S4. ^19^F-1D ligand deconvolutions of enNTS1[Q301C^BTFMA^].......................................................S10

Figure S5. ^19^F-STD pulse scheme............................................................................................................S11

Figure S6. ^19^F-STD decay curves of enNTS1[Q301C^BTFMA^]...................................................................S12

Figure S7. STD decay curves of enNTS1[Q301C^BTFMA^] indicate linear exchange pathway................... S13

Figure S8. [^19^F, ^19^F]-EXSY plot of apo enNTS1[Q301C^BTFMA^]…………...............................................S14

Figure S9. ^19^F-1D transducer deconvolutions of enNTS1[Q301C^BTFMA^]................................................S15

Figure S10. Deconvoluted transducer chemical shift residual error analysis...........................................S16

**SI Tables**.................................................................................................................................................S17

Table S1. mtsslWizard BTFMA rotamer conformers..............................................................................S17

Table S2. Protease digestion mass-spectrometry of enNTS1[Q301C^BTFMA^]...........................................S18

Table S3. ^19^F-T_2_ relaxation times and calculated LWHH of apo enNTS1[Q301C^BTFMA^]........................S19

Table S4. STD exchange rates of enNTS1[Q301C^BTFMA^]........................................................................S20

Experimental Information

***I. Materials and Methods***

**Reagents:**

The construct used for expression of enNTS1[Q301C^BTFMA^] in *Escherichia coli* BL21(DE3) cells was based on a previously published plasmid.^32^ The TALON affinity resin slurry used in the IMAC purification step was purchased from Fisher Scientific (#PI89965). The BTFMA fluorine probe used in all NMR experiments was purchased from Oakwood Chemical (#429825). The Antagonist SR142948 ligand was purchased from Tocris (#2309). All pre-packed affinity resin columns used in the purification of enNTS1[Q301C^BTFMA^] (IMAC, IEX, SEC) was purchased from GE (#45-000-192, #17-5247-01, #29-0452-69). The “MyOne Streptavidin T1” Dynabeads used for ligand saturation assays was purchased from Thermo Fisher (#65601). The fluorescent ligand was made in house using the dye AF647 from AAT Bioquest (#1833). The Red-NTA dye used for the MST assays was purchased from NanoTemper (#L018). The capillaries used for the MST assays was purchased from NanoTemper (#K025). For the cell-based assays, we used NTS1 plasmids cloned into the pCAGGS vector with an N-terminal hemagglutinin-derived signal sequence (ssHA), a FLAG epitope tag and a flexible linker (MKTIIALSYIFCLVFADYKDDDDKGGSGGGGSGGSSSGGG; the FLAG epitope tag is underlined). For the direct NanoBiT-based β-arrestin assay, the N-terminal ssHA-FLAG NTS1 construct was C-terminally fused with the SmBiT with a flexible linker (GGSGGGGSGGSSSGGVTGYRLFEEIL; the SmBiT is underlined). The Human full-length βArr1 was N-terminally LgBiT-fused with the flexible linker and inserted into the pCAGGS vector (LgBiT-β-arrestin1).^23^

**Expression of enNTS1[Q301C^BTFMA^]:**

The enNTS1[Q301C^BTFMA^] plasmid was transformed into BL21(DE3) *Escherichia coli* cells and plated overnight on LB agar supplemented with 100 µg/mL carbenicillin at 37 °C. Liquid cultures of LB media were supplemented with 100 µg/mL carbenicillin, and seeded with overnight colonies; these cultures were incubated overnight at 37 °C and 220 RPM. One-liter 2xYT media growths were inoculated with 9 mL overnight seed supplemented with 100 µg/mL carbenicillin and 0.3% (w/v) glucose. Inoculated cultures were incubated at 37 °C and 220 RPM for approximately 1.25 hours to an OD_600_ of 0.15. The growths were then chilled at 16 °C and 220 RPM to an OD_600_ of 0.6. Finally, each one-liter culture was induced with 0.3 mM IPTG and incubated overnight at 16 °C and 220 RPM for 21 hours. These cultures were harvested via centrifugation at 4,000 g and stored at -80 °C.

^1^**^9^F-labelling and purification of enNTS1:**

A glass beaker was placed on ice and included 50 mL *solubilization buffer* (100 mM HEPES, 400 mM NaCl, 20% (v/v) glycerol, 10 mM MgCl_2_, 10 mM imidazole, pH 8.0), 100 mg lysozyme, 1-unit DNAse, 0.2 mM PMSF, and one protease inhibitor cocktail tablet. Once in solution, cell pellet was added to the beaker and solubilized using a stir bar. Solution was then sonicated on ice: two minutes processing time (10 s on, 20 s off) at 30 AMP. Following sonication, another 0.2 mM PMSF was added to the solution as well as 5 mM ^19^F-BTFMA probe. Solution was stirred at 4 °C for one hour. After incubation, 16 mg aldrithiol was added and the solution incubated for an additional 10 minutes. To remove excess BTFMA probe, a membrane-prep was performed via ultracentrifugation; receptor sample was centrifuged at 100,000 g for 10 minutes. The supernatant was decanted and the same volume of *solubilization buffer* was replaced. This ultracentrifugation step was repeated twice to ensure complete removal of excess BTFMA probe. Following ultracentrifugation, the receptor sample was solubilized with 0.6 mg CHAPS, 0.12 mg CHS, and 15 mL 10% (w/v) DM detergent. The solution was stirred at 4 °C for two hours. After incubation, the *Escherichia coli* lipids were removed via centrifugation at 100,000 g for 10 minutes. The remaining enNTS1 supernatant was then incubated with 12 mL *equilibrated TALON* resin (25 mM HEPES, 10% (v/v) glycerol, 300 mM NaCl, 0.15% (w/v) DM, pH 8.00) at 4 °C for 15 minutes. Following TALON resin binding, the receptor solution was placed into a 50 mL glass gravity column to remove unbound proteins. The TALON resin was then subjected to two subsequent 50 mL wash steps: *TALON wash #1* (25 mM HEPES, 10% (v/v) glycerol, 500 mM NaCl, 0.15% (w/v) DM, 10 mM Imidazole, 4 mM ATP, 10 mM MgCl2, pH 8.0), *TALON wash #2* (25 mM HEPES, 10% (v/v) glycerol, 350 mM NaCl, 0.1% (w/v) LMNG, 10 mM Imidazole, pH 8.0). It is important to note that the second wash step also serves as a detergent exchange step from DM to LMNG. Following detergent exchange, enNTS1 was eluted from the TALON resin via 25 mL *TALON elute buffer* (25 mM HEPES, 10% (v/v) glycerol, 500 mM NaCl, 0.01% (w/v) LMNG, 350 mM Imidazole, pH 8.0) and incubated overnight, or two hours minimum, at 4 °C with 3 mg 3C precision protease. Once the expression proteins (MBP, muGFP) were cleaved from enNTS1, the sample was concentrated to 6 mL in a 50 MWCO concentrator via centrifugation at 3,500 g. To this 6 mL of receptor solution, 60 mL *SP equilibration buffer* (20 mM HEPES, 10% (v/v) glycerol, 0.01% (w/v) LMNG, pH 7.4) was added. This entire solution was then flowed over an equilibrated 5 mL SP ion-exchange column via GE AKTA Pure system (#29018224). The loaded column was then washed with 50 mL *SP wash buffer* (20 mM HEPES, 10% (v/v) glycerol, 250 mM NaCl, 0.01% (w/v) LMNG, pH 7.4). An equilibrated 1 mL Nickel column was attached tandem to the 5 mL SP IEX column, and the receptor eluted via 30 mL *SP elute buffer* (20 mM HEPES, 10% (v/v) glycerol, 1 M NaCl, 0.01% (w/v) LMNG, 25 mM Imidazole, pH 7.4). The enNTS1 solution was then concentrated to 800 µL in a 50 MW concentrator via centrifugation at 3,500 g. The sample was then injected onto a GE S200 size-exclusion chromatography column and flowed at 0.5 mL/min via GE AKTA Pure system and collected in 0.5 mL fractions in *NMR buffer* (20 mM HEPES, 50 mM NaCl, 0.01% (w/v) LMNG, 50 µM TFA, pH 7.5). Following SEC, the desired enNTS1 fractions were pooled, concentrated to 100-300 µM, and flash-frozen via liquid nitrogen and stored at -80 °C. All ligands were added to receptor at 10 Meq prior to NMR data collection. Transducer spectra included 5 Meq transducer, 10 Meq NT8-13, and 10 Meq PIP_2_ in the βArr1 samples. All NMR samples were supplemented with 10% (v/v) D_2_O. Final detergent concentrations for each sample were calculated on known standards of LMNG via ^1^H-1D NMR; all sample LMNG concentrations ranged between 0.01%-0.04% (w/v).

**^19^F-NMR data collection:**

NMR spectra were collected on either a Bruker AVANCE NEO II 14.1 T (Indiana University – Bloomington) or Bruker AVANCE HD 14.1 T (Indiana University School of Medicine) spectrometer equipped with a HCN cryogenic probe tunable to the fluorine frequency. All spectra were collected at 35 °C in 3 mm O.D. tubes. Free induction decay (FID) signals were collected by applying a π/2 pulse length of 13.5 µs, a recycling time of 0.8 ms, and an acquisition time of 0.15 s. A total of 8192 scans were collected, in triplicate, generating a FID comprised of 2499 complex points which were linear predicted to 8,000 points, and apodized with a 30 Hz exponential filter. Triplicate 1Ds were signal averaged to generate the final NMR spectrum. STD, CEST, and T_2_ experiments were collected in a fully-interleaved fashion to account for protein degradation effects on peak intensities/height. 1D saturation transfer experiments were recorded with an acquisition time of 0.12 s, a recycling time of 0.8 ms, and excitation pulse lengths between 50-1000 ms. All saturation pulses had an excitation bandwidth of 8 Hz. T_2_ experiments were collected using a train of 1 ms CPMG spin-echo periods over a 1-6 ms total delay, with an acquisition time of 0.24 s. EXSY experiments were collected with a 100 ms mixing time and an acquisition time of 0.24 s.

**^19^F-NMR spectral deconvolution:**

Processed enNTS1[Q301C^BTFMA^] NMR spectra were deconvoluted using MestReNova. As detailed in a previously published article using this software for ^19^F-NMR analysis on GPCRs, manipulation of individual substate chemical shifts while tracking changes in residual error was performed.^24^ Residual error is defined as the difference between the experimental and simulated resonance, squared, averaged over the region of interest. This procedure was used for every substate analysis discussed in the manuscript to ensure the most accurate deconvolution of NMR spectra.

**Mass spectrometry:**

*Intact protein analysis* - Samples were analyzed on a Synapt G2S equipped with an iClass Acquity HPLC (Waters). Buffer A was 0.1% (v/v) formic acid in water and Buffer B was 0.1% (v/v) formic acid in acetonitrile. Proteins were separated using a nine-minute gradient from 5-99% Buffer B at a flow rate of 50 nL/min. Proteins were separated using a 5 cm x 0.5 mm column in-house packed with Jupiter 5μm C4 resin (Phenomenex). The ToF was set to scan from 400 to 2000 Da at a scan time of 1 second and an analyzer setting of “Resolution”. *Protein processing* - Samples were resuspended and denatured in 8 M urea with 100 mM ammonium bicarbonate (pH 7.8). Disulfide bonds were reduced by incubation for 45 min at 57 °C with a final concentration of 10 mM Tris (2-carboxyethyl) phosphine hydrochloride (#C4706, Sigma Aldrich). A final concentration of 20 mM iodoacetamide (#I6125, Sigma Aldrich) was then added to alkylate these side chains and the reaction was allowed to proceed for one hour in the dark at 21 °C. Samples were diluted to 1 M urea using 100 mM ammonium bicarbonate, pH 7.8. Trypsin (V5113, Promega) or chymotrypsin (#11418467001, Sigma Aldrich) was added at a 1:100 ratio and the samples were digested for 14 hours at 37 °C. *Mass spectrometry* - individual samples were desalted using ZipTip pipette tips (EMD Millipore), dried down and resuspended in 0.1% (v/v) formic acid. Fractions were analyzed by LC-MS on an Orbitrap Fusion Lumos equipped with an Easy NanoLC1200 HPLC (Thermo Fisher Scientific). Buffer A was 0.1% (v/v) formic acid in water. Buffer B was 0.1% (v/v) formic acid in 80% acetonitrile. Peptides were separated on a 30-minute gradient from 0-3% Buffer B. Precursor ions were measured in the Orbitrap with a resolution of 120,000. Fragment ions were measured in the Orbitrap with a resolution of 15,000. The spray voltage was set at 1.8 kV. Orbitrap MS1 spectra (AGC 1×10^6^) were acquired from 350-2000 m/z followed by data-dependent HCD MS/MS (collision energy 30%, isolation window of 2 Da) for a 3 s cycle time. Charge state screening was enabled to reject unassigned and singly charged ions. A dynamic exclusion time of 30 s was used to discriminate against previously selected ions. *Database search* - The LC-MS/MS data was searched against the protein sequence using Protein Prospector (v5.22.1). The database search parameters for the tryptic search allowed for two missed cleavages and one non-tryptic cleavage. The search parameters for the chymotryptic search allowed for four missed cleavages and one non-chymotryptic cleavage. A precursor and fragment mass tolerance of 10 ppm was used. Oxidation of methionine, pyroglutamine on peptide amino termini, carbamidomethylation of cysteine, and protein N-terminal acetylation were set as variable modifications. In addition, modification of cysteine residues by BTFMA (C_9_H_6_F_3_NO) was set as a variable modification.

**Dynabead ligand saturation assay:**

Receptor purifications ultimately used for experimental data collection were tested for proper transmembrane folding via agonist NT8-13 saturation binding assay. The enNTS1 construct mentioned above possesses an N-terminal Avi-tag; this affinity tag was used to immobilize the purified receptor to magnetic Dynabeads that possessed the complementary Streptavidin protein. A Promega (#V8351) 96-well magnetic side strip plate allowed for the capture of the Dynabead-enNTS1[Q301C^BTFMA^] complex. This complex (25 nM) was subjected to increasing amounts of fluorescent endogenous ligand NT8-13^AF647^ (0.5-500 nM), made in-house. Unbound fluorescent peptide was washed away and resultant Dynabead-receptor-ligand complex fluorescence intensity measured via plate reader (BioTek Synergy Neo2). This experimental setup was also used to assay functional stability of enNTS1[Q301C^BTFMA^] incubated at 37 °C.

**Gα_q_ peptide pulldown assay:**

This assay makes use of biotinylated G protein peptides and magnetic Dynabeads. For this assay, an Avi-tag-less enNTS1 construct with a C-terminal muGFP reporter protein was generated. All experimental procedures followed are previously published using a N-terminal biotinylated (bioQp) peptide corresponding to residues 333-359 of the Gα_q_ C-terminus (a.k.a. α5-helix).^26^ Percentages of enNTS1 pulled-down via Gα_q_ peptide were assessed in varying liganded conditions. enNTS1 (30 nM) was incubated with 10 Meq Gα_q_ peptide and ligand (30 µM). Complexed receptor was magnetically pulled down and separated from unbound receptor. Unbound receptor was washed away and resultant Dynabead-receptor-Gα_q_ peptide complex fluorescence intensity measured via plate reader (BioTek Synergy Neo2). All samples were collected in both technical and instrumental triplicate.

**βArr1[ΔCT] MST assay:**

This assay makes use of NTA-labeled βArr1[ΔCT] (RED-βArr1[ΔCT]) and NanoTemper premium capillaries. For this assay, constant concentrations of RED-βArr1[ΔCT] and ligand were incubated with varying concentrations of enNTS1 and subsequently assayed via NanoTemper Monolith NT MST instrument. Each experiment consisted of 16 samples of 25 nM RED-βArr1[ΔCT], 22.5 µM ligand and PIP2, and a 1:2 dilution series of enNTS1 with a starting concentration of 2.25 µM. Data was collected using 75% LED and 80% MST power, then extracted via “T-Jump” analysis from NanoTemper analysis software. Data was normalized between the maximum signal and average of four smallest signals. Data was fit to the quadratic binding function. All samples were collected in both technical and instrumental triplicate.

**TGFα shedding assay:**

The TGFα shedding assay was performed as described previously.^22^ HEK293A cells (Thermo Fisher Scientific) were seeded in a 6-well culture plate (Greiner Bio-One) at a concentration of 2x10^5^ cells/ml (2 ml per well hereafter) in DMEM (Nissui Pharmaceutical) supplemented with 10% (v/v) FBS (Gibco), glutamine, penicillin, and streptomycin, one day before transfection. The transfection solution was prepared by combining 5 µl of 1 mg/ml polyethylenimine Max solution (Polysciences) and a plasmid mixture consisting of 200 ng ssHA-FLAG-NTS1 and 500 ng alkaline phosphatase (AP)-tagged TGF-α (AP-TGFα; human codon-optimized). One-day after incubation, the transfected cells were harvested by trypsinization, neutralized with DMEM containing 10% (v/v) FCS and penicillin–streptomycin, washed once with Hank’s Balanced Salt Solution (HBSS) containing 5 mM HEPES (pH 7.4), and resuspended in 6 ml of the HEPES-containing HBSS. The cell suspension was seeded into a 96-well plate at a volume of 80 µl (per well hereafter) and incubated for 30 minutes in a CO_2_ incubator. A test ligand (diluted in 0.01% (w/v) BSA and 5 mM HEPES-containing HBSS at 10x concentration) or vehicle was added at a volume of 10 µl. After 5 min, a test agonist (NT8-13), (serially diluted in 0.01% (w/v) BSA and 5 mM HEPES-containing HBSS at 10x concentration) were added and the plate was incubated for 1-hour. After centrifugation, conditioned media (80 µl) was transferred to an empty 96-well plate. AP reaction solution (10 mM p-nitrophenylphosphate (*p*-NPP), 120 mM Tris–HCl (pH 9.5), 40 mM NaCl, 10 mM MgCl_2_) was dispensed into the cell culture plates and plates containing conditioned media (80 µl). Absorbance at 405 nm was measured before and after a 1-hour or 2-hour incubation at room temperature using a microplate reader (SpectraMax 340 PC384; Molecular Devices). Ligand-induced AP-TGF-α release was calculated as described previously.^22^ Unless otherwise noted, vehicle-treated AP-TGF-α release signal was set as a baseline. Using Prism 8 software (GraphPad Prism), AP-TGF-α release signals were fitted with a four-parameter sigmoidal concentration-response curve.

**NanoBiT-based βArr1 assay:**

The NanoBiT-based βArr1 assay was performed as described previously.^23^ HEK293A cells were seeded in a 6-cm culture dish (Greiner Bio-One) at a concentration of 2x10^5^ cells/ml (4 ml per dish) in the FBS-supplemented DMEM. Plasmid transfection was performed by combining 10 µl of the polyethylenimine Max solution and a plasmid mixture consisting of 1 µg ssHA-FLAG-GPCR-SmBiT and 200 ng LgBiT-βArr1 in 400 µl of Opti-MEM. After incubation for one day, the transfected cells were harvested with 0.5 mM EDTA-containing Dulbecco’s PBS (D-PBS), centrifuged, and suspended in 4 ml of HBSS containing 0.01% (w/v) BSA and 5 mM HEPES (pH 7.4) (assay buffer). The cell suspension was dispensed in a white 96-well plate (Greiner Bio-One) at a volume of 70 µl per well and loaded with 20 µl of 50 µM coelenterazine (Carbosynth), diluted in the assay buffer. After a 2-hour incubation at room temperature, the plate was measured for its baseline luminescence (SpectraMax L, 2PMT model, Molecular Devices). Thereafter, a test ligand (diluted in the assay buffer at 10x concentration) was added at a volume of 10 µl and the plate was incubated for 15 min at room temperature. After a second measurement of luminescence, a test agonist (NT8-13), (serially diluted in the assay buffer at 6x concentration) were added at a volume of 20 µl and the plate was immediately read as a kinetics mode for 10 min. Luminescence counts recorded from 5 min to 10 min after the agonist addition were averaged and normalized to the initial counts. The fold-change signals were further normalized to the vehicle-treated signal and were plotted as a βArr1 recruitment response. Using the Prism 8 software, the βArr1 recruitment signals were fitted to a four-parameter sigmoidal concentration-response curve.

**Flow cytometry analysis:**

Plasmid transfection into HEK293A cells were performed as described in the TGFα shedding assay section. One-day after transfection, the cells were collected by adding 200 μl of 0.53 mM EDTA-containing D-PBS, followed by 200 μl of 5 mM HEPES (pH 7.4)-containing HBSS. The cell suspension was transferred to a 96-well V-bottom plate in duplicate, blocked with 2% (v/v) goat serum- and 2 mM EDTA-containing D-PBS (blocking buffer; 100 µL per well hereafter) and fluorescently labeled with the anti-FLAG-epitope tag monoclonal antibody (Clone 1E6, FujiFilm Wako Pure Chemicals; 10 µg per ml in the blocking buffer; 25 µL) and a goat anti-mouse IgG secondary antibody conjugated with Alexa Fluor 488 (Thermo Fisher Scientific, 10 μg per ml diluted in the blocking buffer; 25 µL). Live cells were gated with a forward scatter (FS-Peak-Lin) cutoff at the 390 setting, with a gain value of 1.7 and fluorescent signal derived from Alexa Fluor 488 was recorded in the FL1 channel.

SI Figures

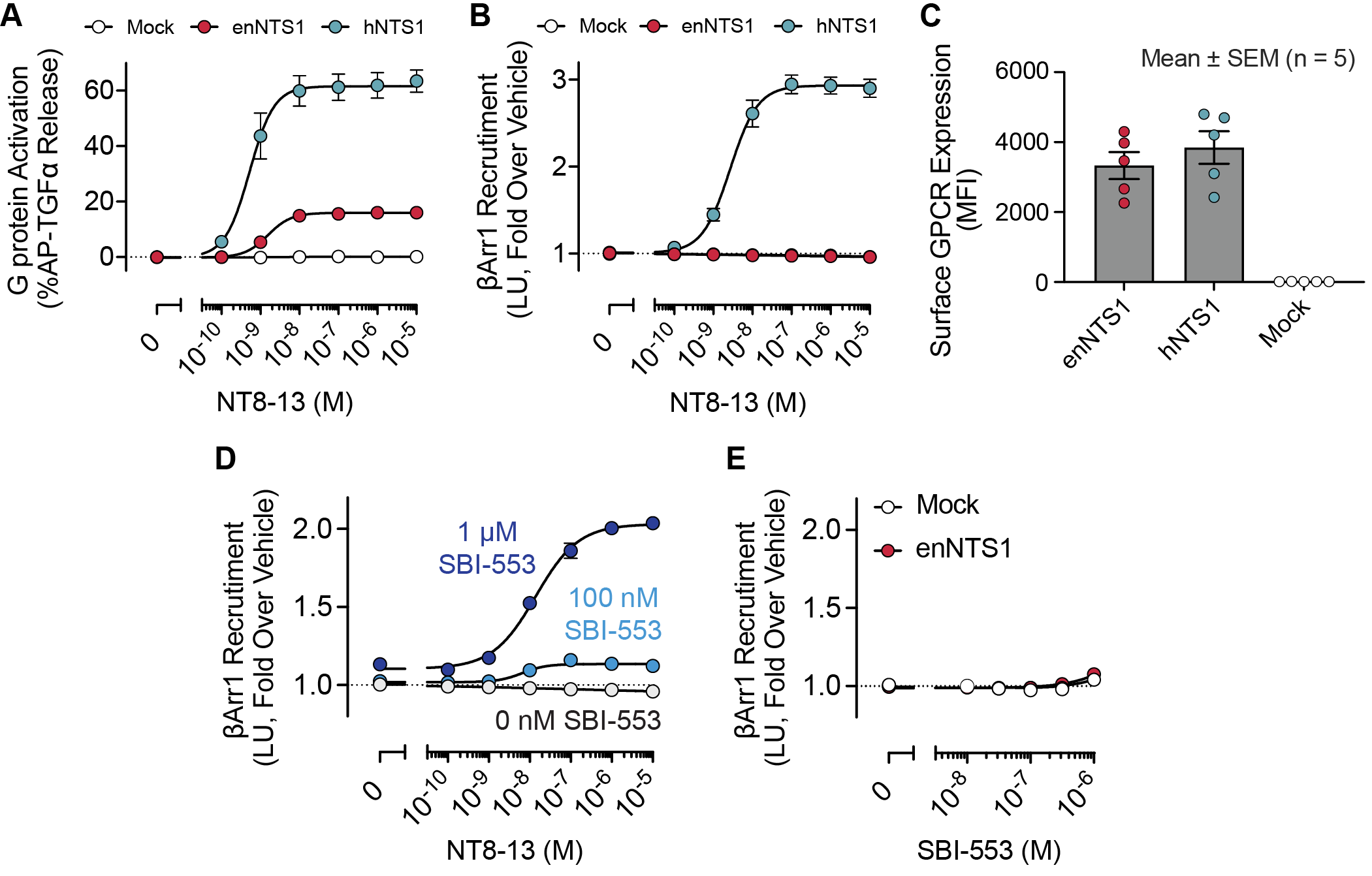

**Figure S1.** G protein activation and βArr1 recruitment cellular assays. Dose-dependent (A) TGFα shedding and (B) βArr1 recruitment assays for HEK293A cells transiently-transfected with enNTS1, hNTS1, or V2R mock. Error bars represent SEM from three independent experiments. (C) Surface expression levels for each receptor were measured using flow cytometry. Error bars represent SEM from five independent experiments. (D) Addition of 0, 100 nM, or 1 μM βArr1-biased allosteric modulator (SBI-553) potentiates NT8-13-dependent βArr1 recruitment. (E) SBI-553 alone is unable to substantially stimulate βArr1 recruitment. Please note for many samples, size of error bar is smaller than symbol.

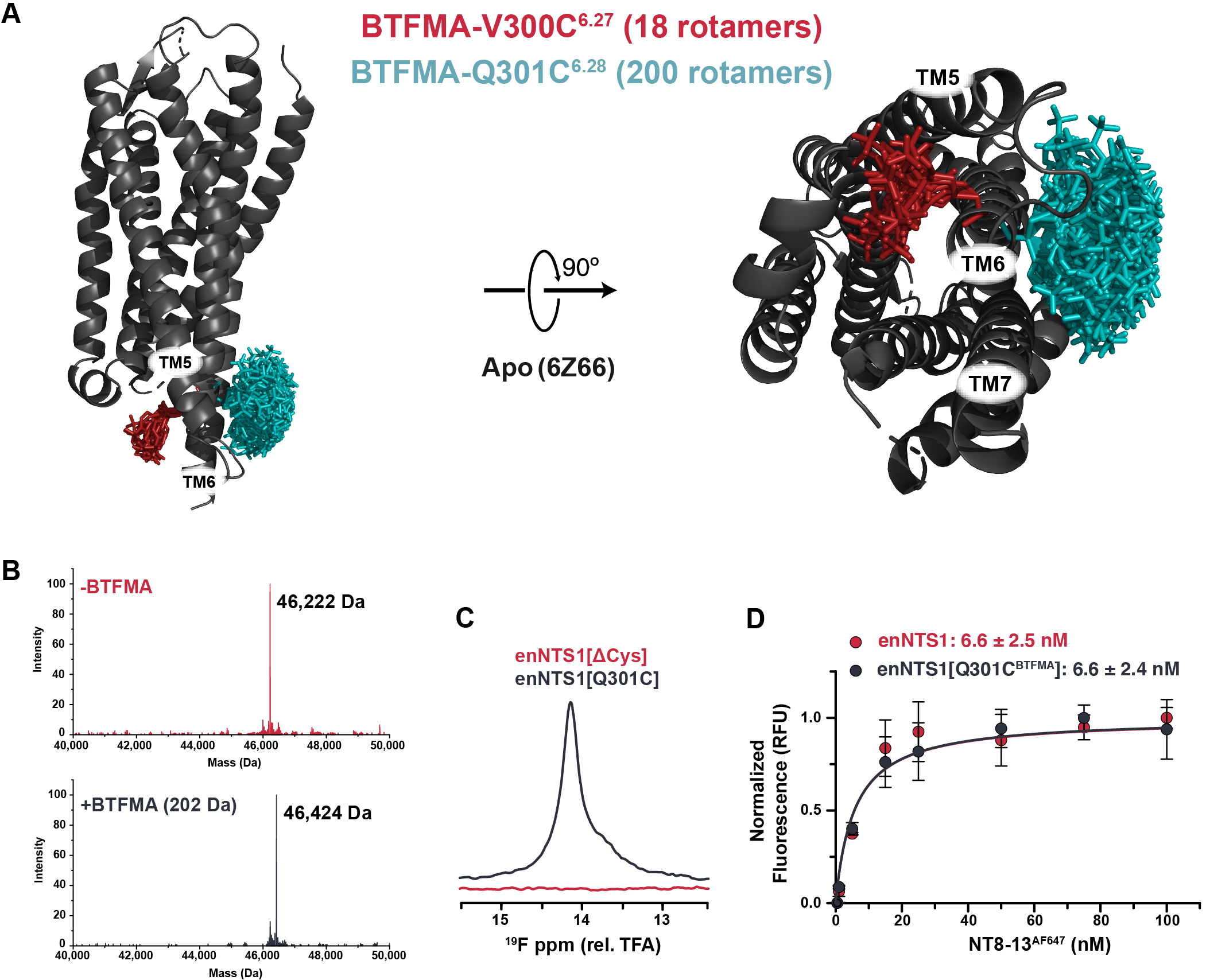

**Figure S2.** Selective ^19^F-labeling of enNTS1 intracellular helix. (A) NTS1 V300C^6.27^ vs Q301C^6.28^ mtsslWizard ^19^F-BTFMA rotamers in apo-state crystal structure (PDB 6Z66). Modeled ^19^F-labeling at residue 6.27 is sterically hindered by tight TM5/TM6 helix packing, generating 18 possible rotamer conformations (red). BTFMA probe mobility is unrestricted at residue 6.28 (teal). (B) Mass spectrometry of intact enNTS1[Q301C] (red) and enNTS1[Q301C^BTFMA^] (slate). A single covalently-attached BTFMA molecule corresponds to a 202 Da mass shift. (C) ^19^F-NMR spectra of enNTS1[Q301C^BTFMA^] (slate) and enNTS1[ΔCys^BTFMA^] (red) where all exposed cysteine residues were mutated to serine; chemical shifts are relative to the resonance of trifluoroacetic acid (TFA). (D) DynaBead saturation binding assay of enNTS1 (red) and enNTS1[Q301C^BTFMA^] (slate) using fluorescent-tagged NT8-13 agonist (NT8-13^AF647^). Data points represent the average normalized RFU from three individual experiments; error bars represent the standard deviation. Biomolecular equilibrium dissociation constant (K_d_) calculated from a global fit of experimental data using the quadratic binding model.

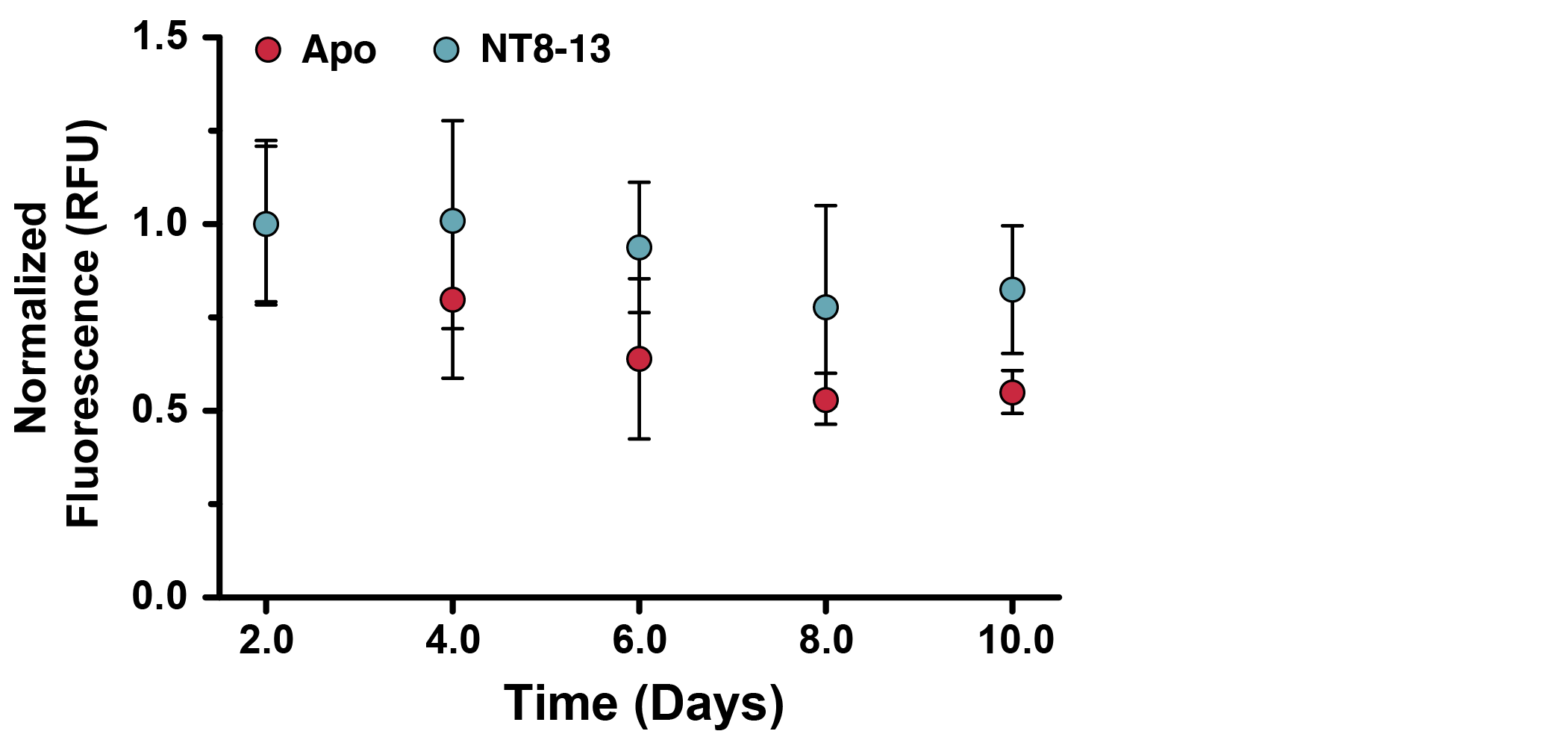

**Figure S3.** Long-term stability of enNTS1[Q301C^BTFMA^]. enNTS1[Q301C^BTFMA^] was incubated at 37 °C for varying amounts of time in the presence (teal) or absence (red) of NT8-13^AF647^. After the indicated time, all samples were incubated with 10 Meq NT8-13^AF647^ and functionally assessed via Dynabead assay. Data points represent the average normalized RFU from six individual experiments collected in instrument triplicate; error bars represent the standard deviation.

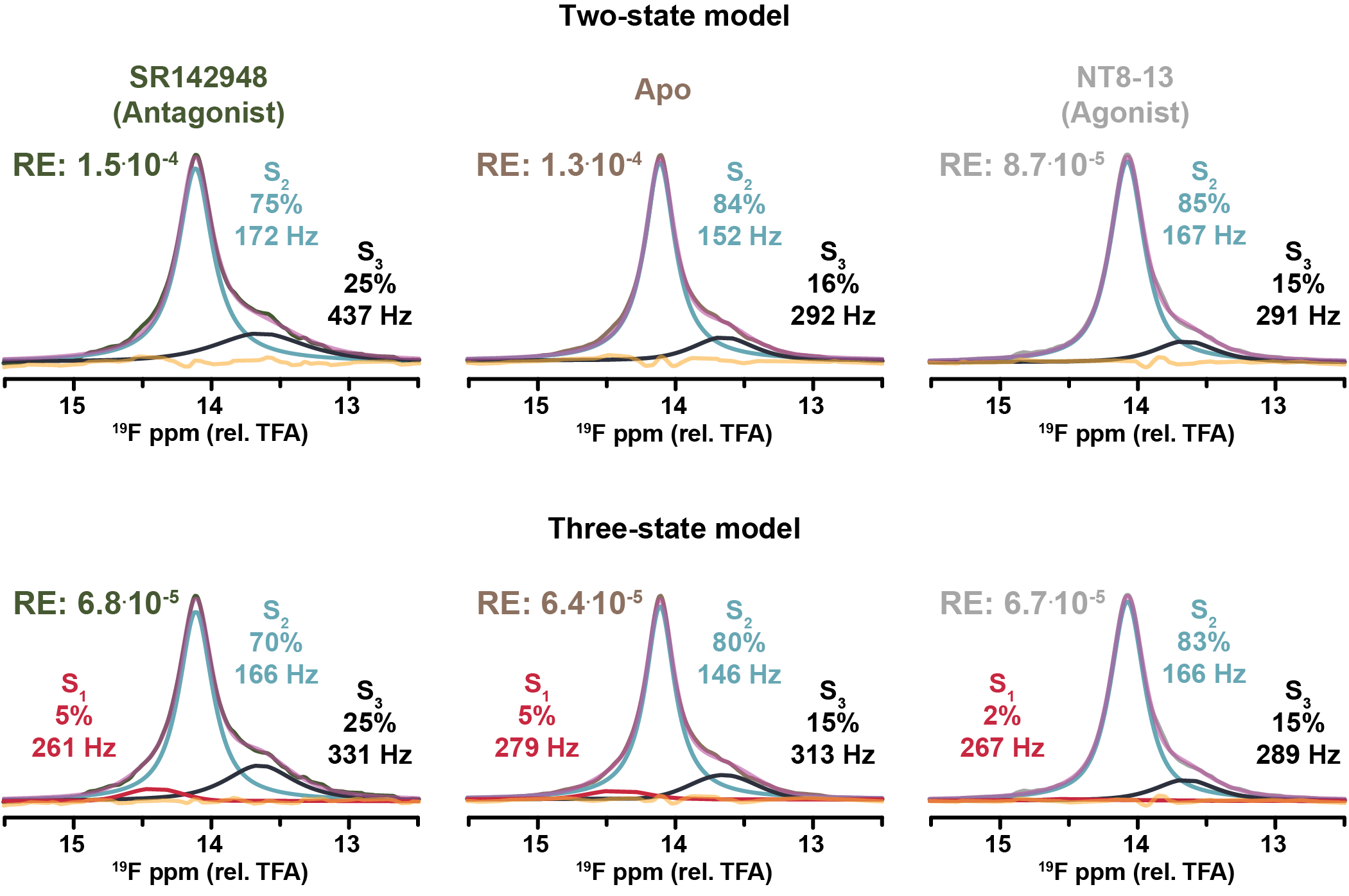

**Figure S4.** Deconvolution of enNTS1[Q301C^BTFMA^] ^19^F-NMR spectra. 1D spectra of apo and ligand-bound enNTS1[Q301C^BTFMA^] samples deconvoluted assuming two or three Lorentzian lineshapes. Deconvoluted resonances were used to simulate the original spectrum (magenta) for each condition. At each frequency, the difference between experimental and simulated spectra is indicated as a residual (orange). The residual error (RE) across the 12.5-15.5 ppm spectral region is the average of residuals, squared, at each frequency.

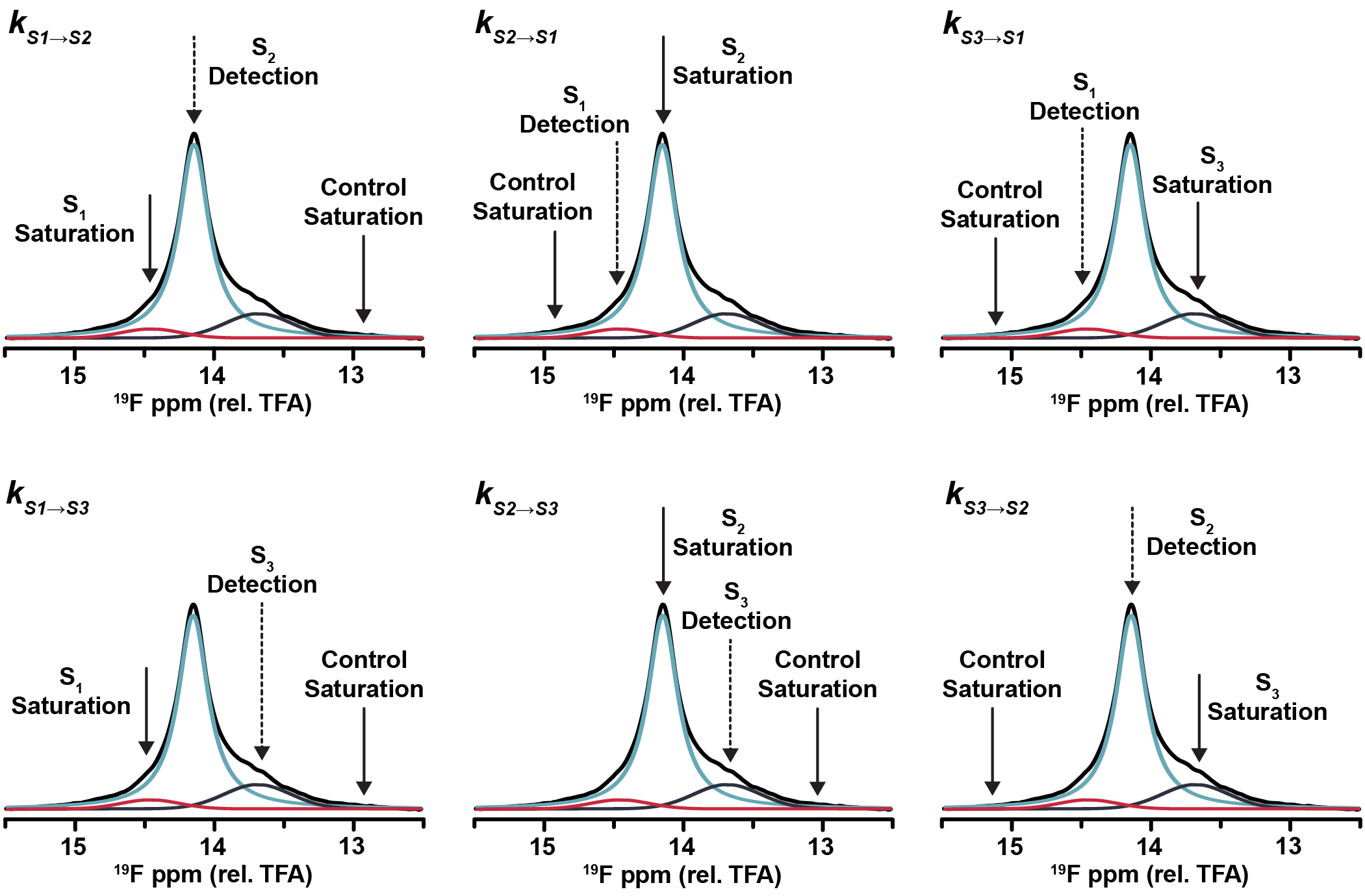

**Figure S5.** ^19^F-STD pulse scheme diagram. Approximate saturation frequencies utilized in the ^19^F-STD measurements of enNTS1[Q301C^BTFMA^]. All saturation pulses had an excitation bandwidth of 8 Hz.

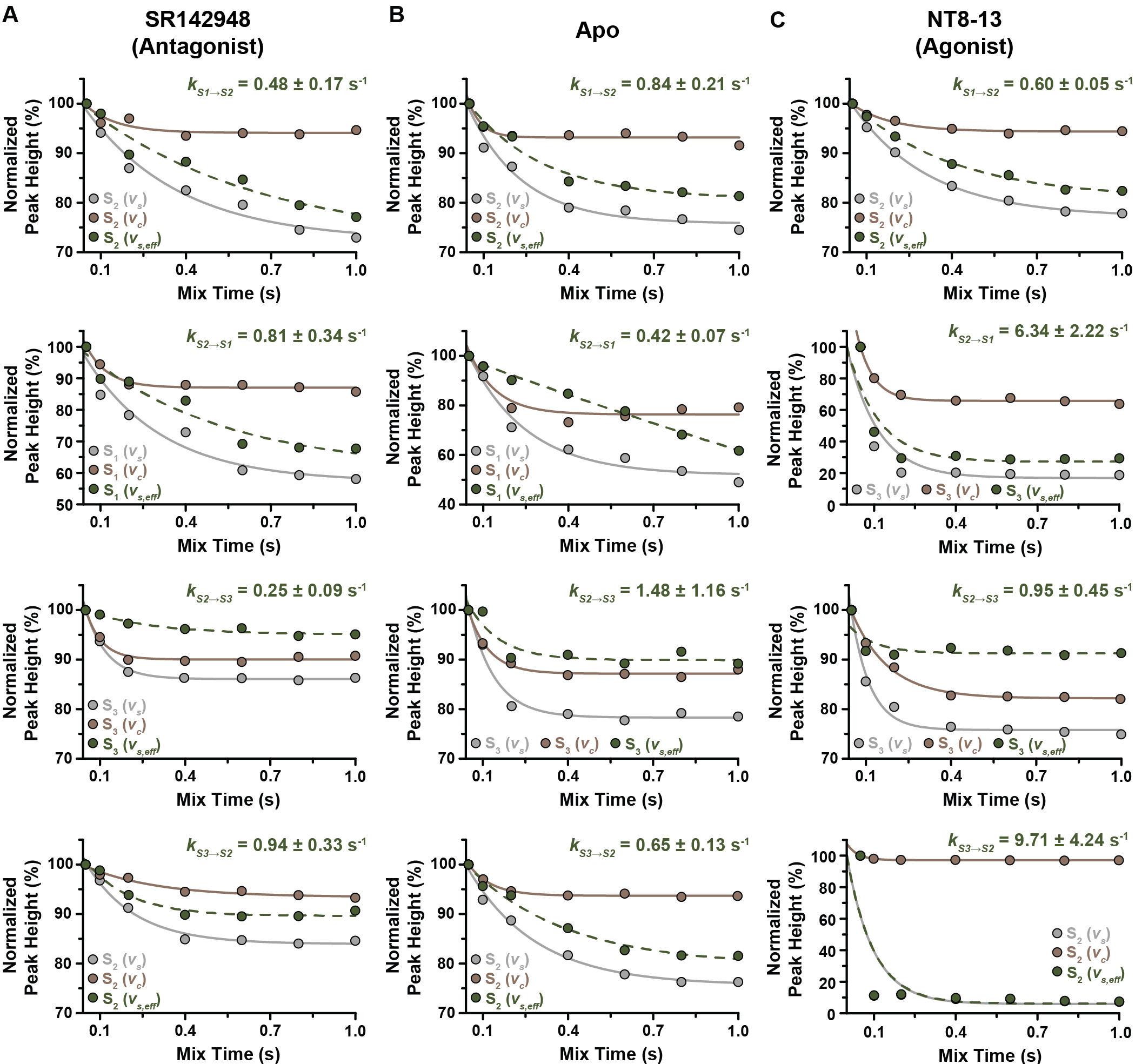

**Figure S6.** ^19^F-STD decay curves of enNTS1[Q301C^BTFMA^]. (A,B,C) STD decay curves from deconvoluted ^19^F-NMR spectra of liganded enNTS1[Q301C^BTFMA^]. Indicated ligands were incubated with receptor at 10 Meq. In each graph, the grey curve represents the peak height following on-resonance saturation of a defined substate (*v_s_*); brown curves represent the control peak height when control saturation (off-resonance) was applied equidistance, but opposite frequency, from the saturation experiment (*v_c_*). The effective decay curve (*v_s,eff_*; dashed green line) is the difference of on- and off-resonance experiments fit to the Bloch-McConnell equations.

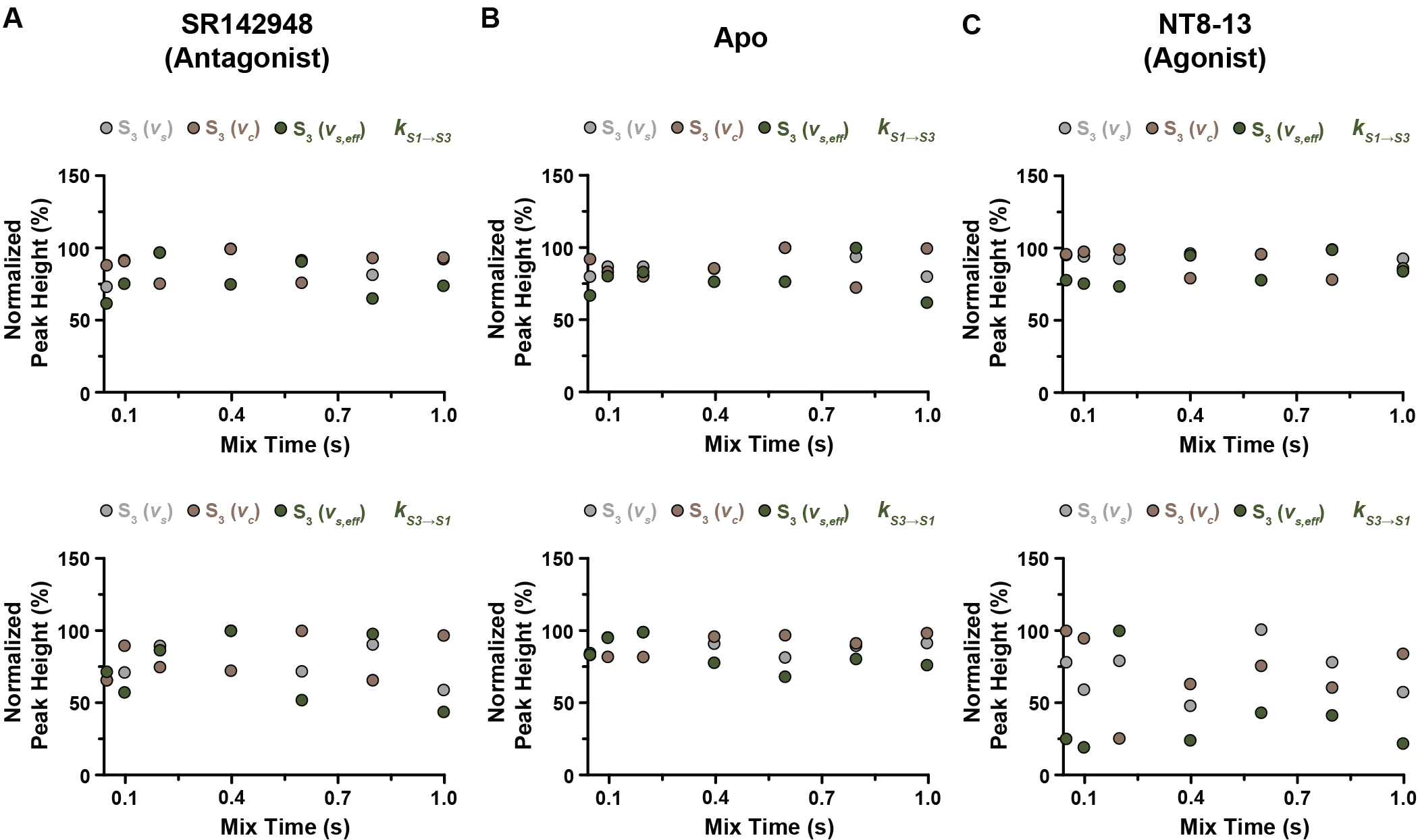

**Figure S7.** STD decay curves of enNTS1[Q301C^BTFMA^] indicate linear exchange pathway. ^19^F-STD curves associated with enNTS1[Q301C^BTFMA^] in the antagonist- (A), apo (B), or agonist-bound (C) states. Ligands incubated with receptor at 10 Meq. In each graph, the grey spheres represent the peak height following on-resonance saturation of a defined substate (*v_s_*); brown spheres represent the control peak height when control saturation (off-resonance) was applied equidistance, but opposite frequency, from the saturation experiment (*v_c_*). The effective decay curve (*v_s,eff_*; green spheres) is the difference of on- and off-resonance experiments. As no curve can be fit for S1🡨🡪S3 transition, the exchange pathway is linear (S1🡨🡪S2🡨🡪S3) not triangular.

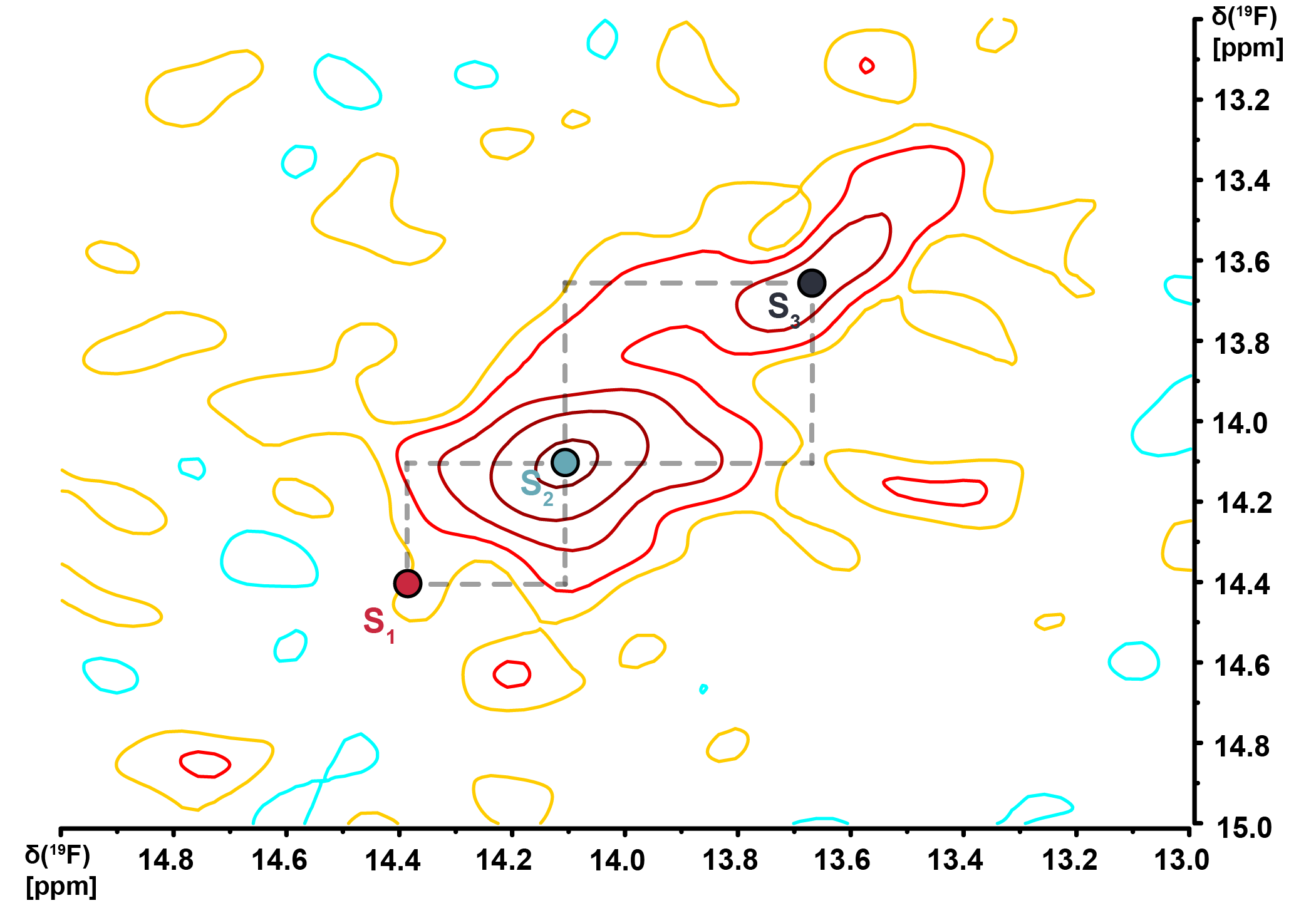

**Figure S8.** [^19^F, ^19^F]-EXSY plot of apo enNTS1[Q301C^BTFMA^]. A 2D [^19^F, ^19^F]-EXSY spectrum of apo enNTS1[Q301C^BTFMA^] labeled with diagonal peak positions of S_1_, S_2_, and S_3_. No exchange cross peaks were observed.

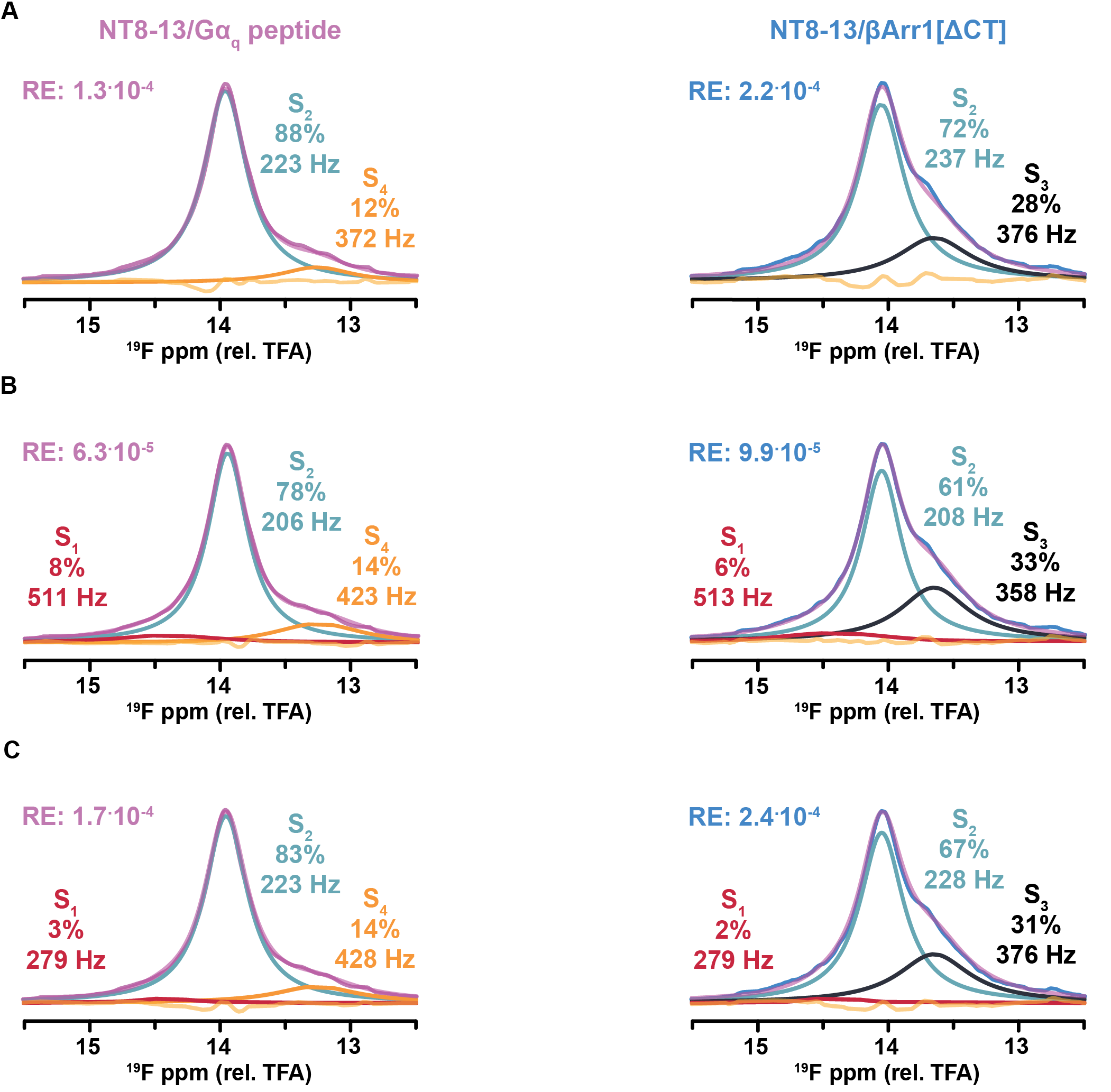

**Figure S9.** ^19^F-1D transducer deconvolutions of enNTS1[Q301C^BTFMA^]. ^19^F-1D spectra of agonist-bound enNTS1[Q301C^BTFMA^] complexed with either Gα_q_ peptide (*left*) or βArr1[ΔCT] (*right*) deconvoluted without S_1_ (A), including S_1_ with an unconstrained LWHH (B), and including S_1_ with a constrained LWHH (C). Deconvoluted resonances were used to simulate the original spectrum (magenta) for each condition. At each frequency, the difference between experimental and simulated spectra is indicated as a residual (orange). The residual error (RE) across the 12.5-15.5 ppm spectral region is the average of residuals, squared, at each frequency.

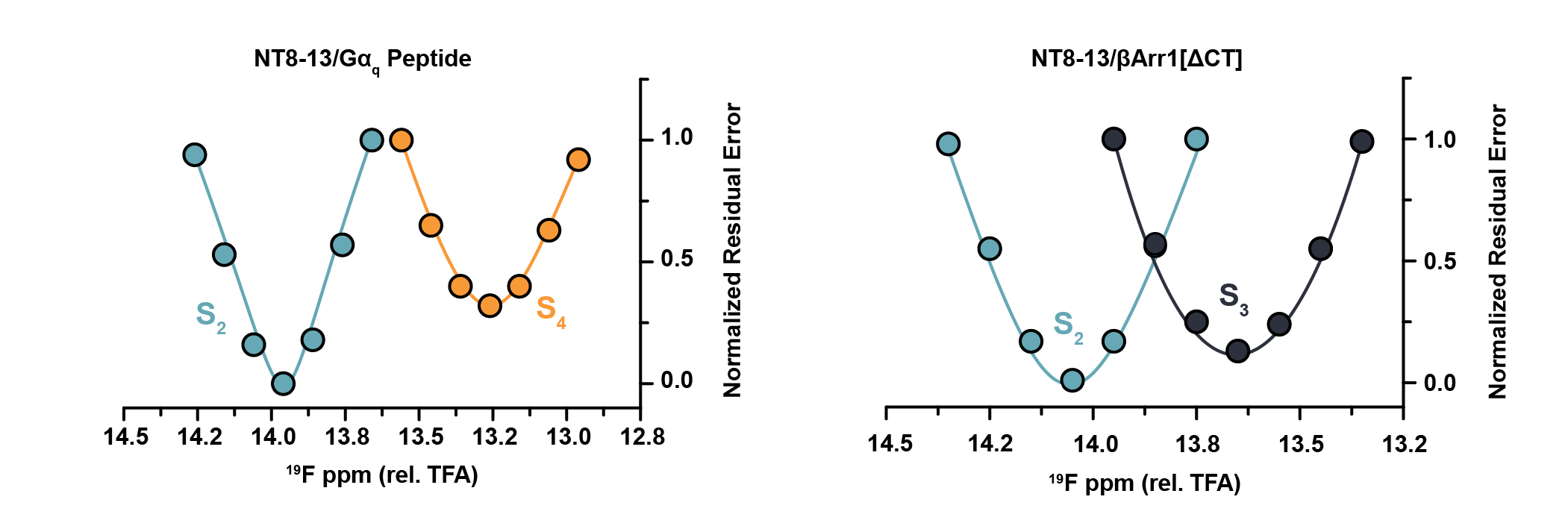

**Figure S10.** Deconvoluted transducer chemical shift residual error analysis. ^19^F-NMR spectra for Gα_q_ peptide and βArr1[ΔCT] ternary complexes were deconvoluted into component substates using MestReNova. LWHH and peak height were constrained while individual substate chemical shifts were modulated and residual error recorded. The lowest residual error value for each substate represents the chemical shift used in deconvolution. Residual error values were normalized prior to graphing.

SI Tables

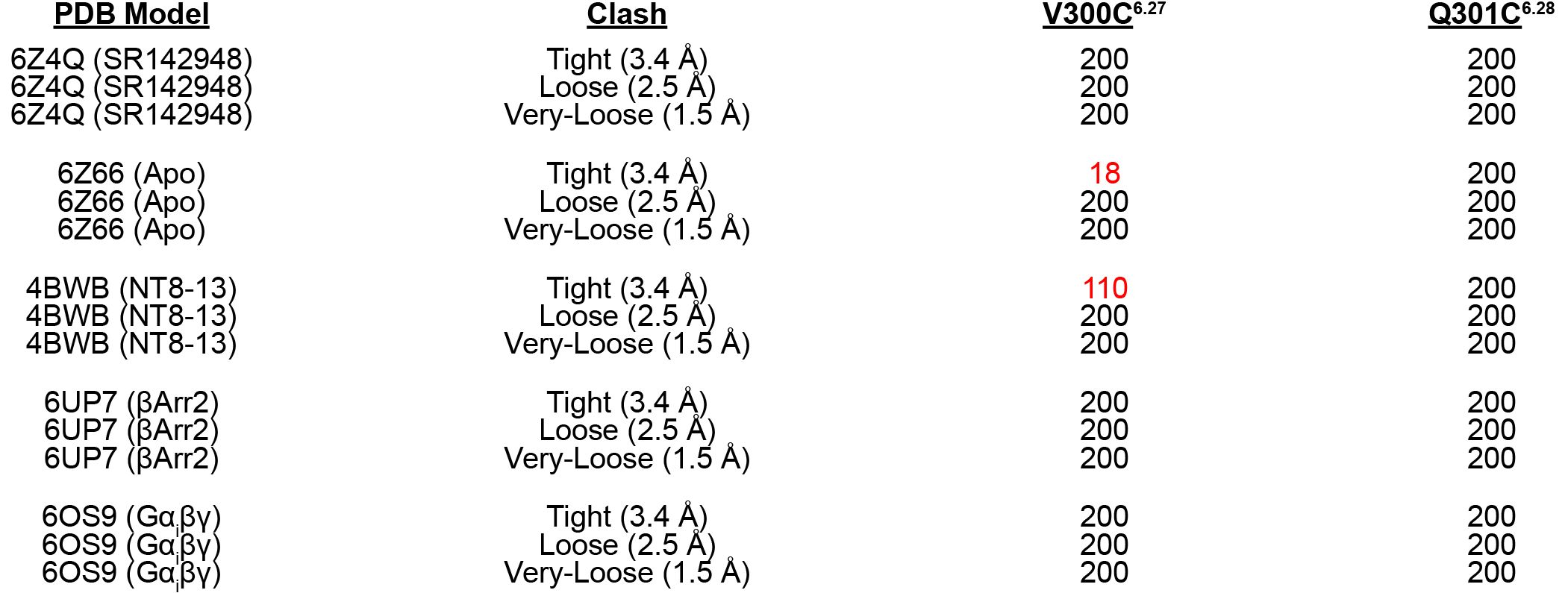

**Table S1.** mtsslWizard BTFMA rotamer conformers. NTS1 PDB models were discretely labeled with BTFMA, at indicated position, in silico via mtsslWizard. A total of 200 rotamer conformers were assessed at varying degrees of tightness. Numbers listed indicate total number of rotamers without steric clash.

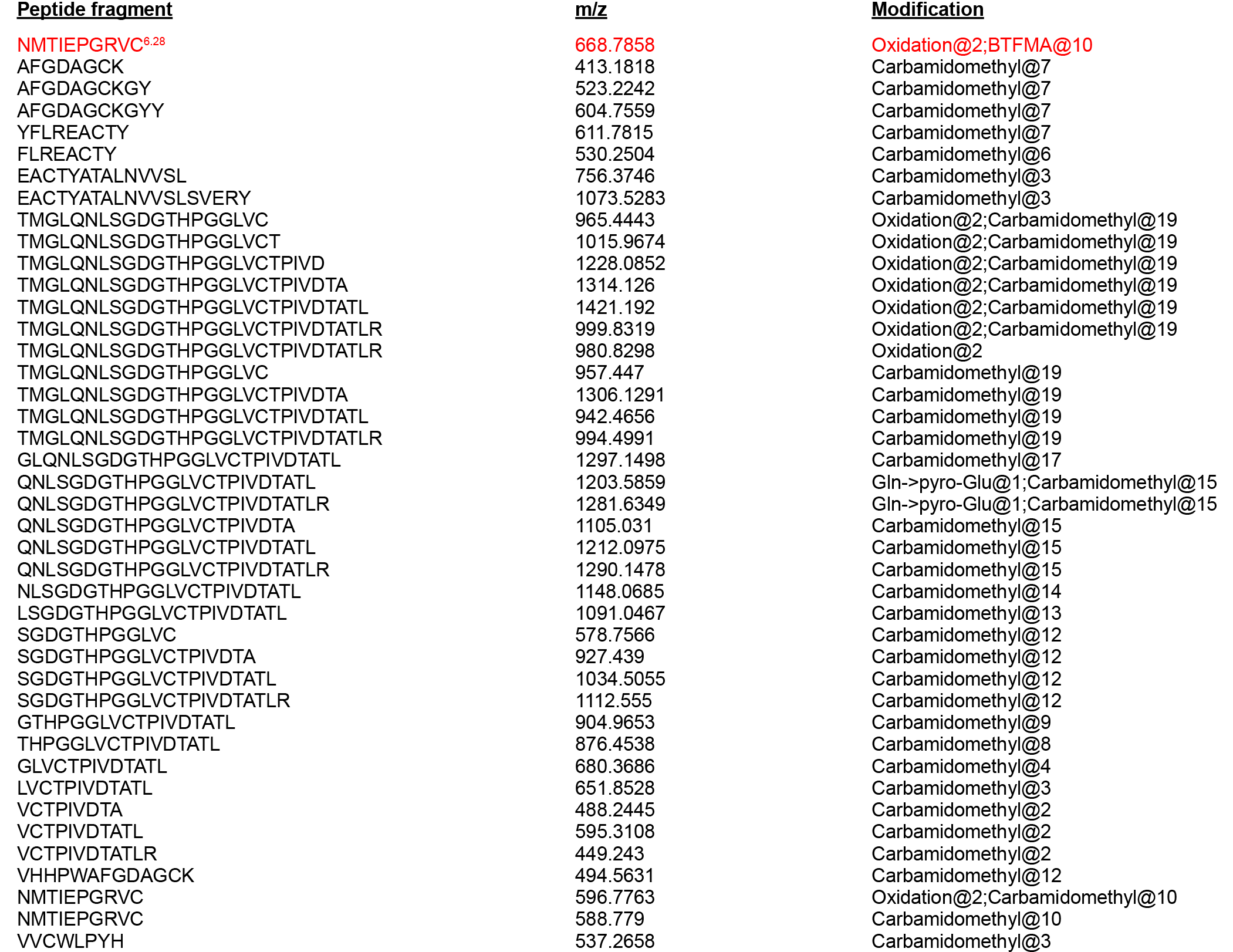

**Table S2.** Protease digestion mass-spectrometry of enNTS1[Q301C^BTFMA^]. Purified enNTS1[Q301C^BTFMA^] digested with trypsin or chymotrypsin, and subsequently analyzed via MS. All five cysteines present in the primary sequence detected through various peptide fragments. Mass over charge (m/z) ratio and residue modification for each peptide fragment listed. Note only C301^6.28^ (red) incorporates BTFMA during labeling.

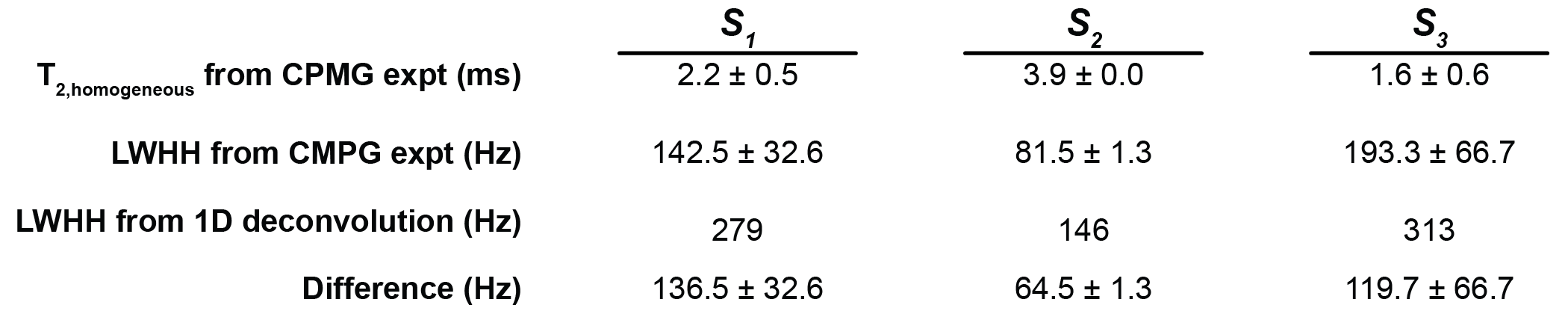

**Table S3.** Analysis of apo enNTS1[Q301C^BTFMA^] T_2_ relaxation times and LWHH. Substate peak heights from CPMG T2 series were fitted to two- or three-parameter mono-exponential functions for direct determination of T_2, homogenous_ and LWHH using the relationship (LWHH = 1/(π*T_2_). 1D ^19^F-spectral deconvolutions simultaneously fitted peak area, chemical shift, and LWHH.

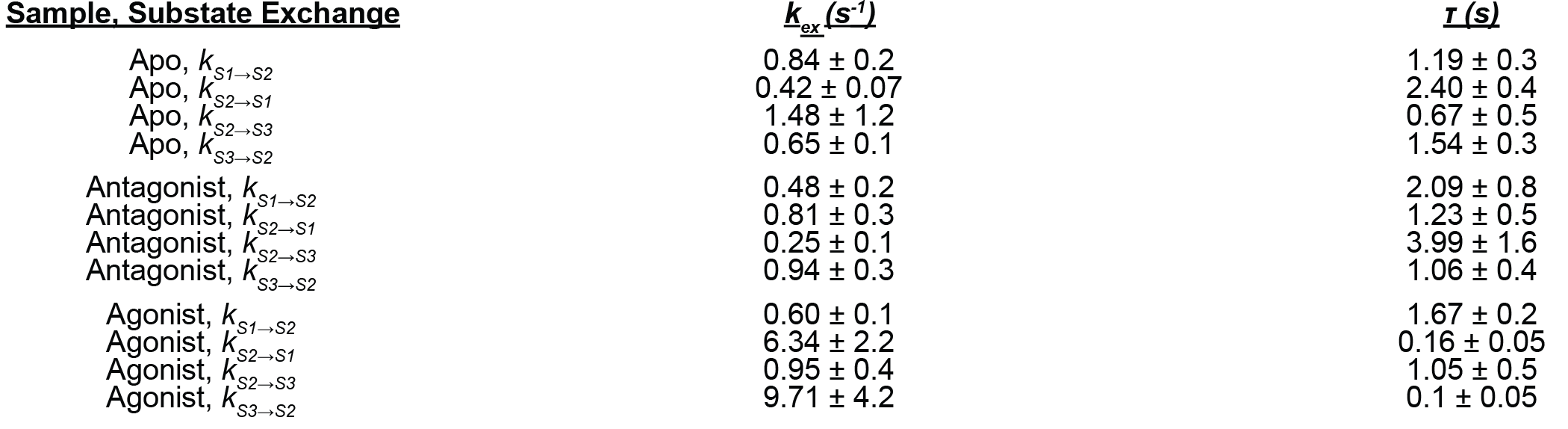

**Table S4.** STD exchange rates of enNTS1[Q301C^BTFMA^]. Exchange rates from deconvoluted ^19^F-NMR STD spectra of enNTS1[Q301C^BTFMA^]. Receptor was incubated with 10 Meq ligand. Rates calculated from fitting effective (*v*_s,eff_) to Bloch-McConnell equations.
